# Guided nuclear exploration increases CTCF target search efficiency

**DOI:** 10.1101/495457

**Authors:** Anders S. Hansen, Assaf Amitai, Claudia Cattoglio, Robert Tjian, Xavier Darzacq

## Abstract

Mammalian genomes are enormous. For a DNA-binding protein, this means that the number of non-specific, off-target sites vastly exceeds the number of specific, cognate sites. How mammalian DNA-binding proteins overcome this challenge to efficiently locate their target sites is not known. Here through live-cell single-molecule tracking, we show that CCCTC-binding factor, CTCF, is repeatedly trapped in small zones in the nucleus in a manner that is largely dependent on its RNA-binding region (RBR). Integrating theory, we devise a new model, Anisotropic Diffusion through transient Trapping in Zones (ADTZ), to explain this. Functionally, transient RBR-mediated trapping increases the efficiency of CTCF target search by ∼2.5 fold. Since the RBR-domain also mediates CTCF clustering, our results suggest a “guided” mechanism where CTCF clusters concentrate diffusing CTCF proteins near cognate binding sites, thus increasing the local ON-rate. We suggest that local “guiding” may represent a general target search mechanism in mammalian cells.

## INTRODUCTION

Mammalian nuclei are organized by a myriad of biophysical forces into sub-compartments with distinct functions. At the micron-scale, nuclear compartments such as nucleoli, speckles, and Cajal bodies, carry out specialized biochemical functions that are spatially segregated (Mao et al., 2011). Below the micron-scale and diffraction-limit of conventional optical microscopy, many proteins interact dynamically to form local high concentration clusters or hubs (Banjade and Rosen, 2014; Boehning et al., 2018; Chong et al., 2018). Thus, many proteins exhibit an organized and non-random nuclear distribution. At the same time, several proteins display anomalous and non-Brownian diffusion inside the nucleus (Höfling and Franosch, 2013; Metzler et al., 2014; Rhodes et al., 2017), which has been proposed to be due to molecular crowding and transient interactions (Bancaud et al., 2012; Woringer and Darzacq, 2018). It is still unclear what mechanisms allow proteins to form stable or transient clusters and control their diffusion and target search mechanism *in vivo*. It has been proposed that some proteins with disordered domains are capable of forming hubs that behave as distinct liquid droplets or condensates (Feric et al., 2016). In such structures, protein-RNA interactions have been shown to be important (Lin et al., 2015). The most obvious example is the nucleolus (Weber and Brangwynne, 2015), which is rich in ribosomal RNA. However, the functional role of these structures in loading, guiding and tightly regulating the functional search kinetics of associated proteins has yet to be elucidated *in vivo*.

Since the kinetics of a reaction or binding event depend on the diffusive properties and nuclear organization of the reactants (Bénichou et al., 2010; Rice, 1985), understanding the molecular interactions that control a nuclear protein’s target search mechanism and its overall distribution is essential to understanding its function. Target search mechanisms have been extensively probed in prokaryotes (Hammar et al., 2012; Kapanidis et al., 2018), where current models (Bauer and Metzler, 2012) suggest that proteins find their target sites through facilitated diffusion (Slutsky and Mirny, 2004). In this family of models, proteins are suggested to move according to two principal modes: (1) sliding in 1D along the DNA strand; (2) occasionally disassociating from the strand, diffusing in 3D, only to bind at a proximal DNA site. It is thought that a mixture of these two modes of motion, with specific rates of transition between the states, would serve as an optimal search strategy (Lomholt et al., 2005). However, how DNA-binding proteins find their targets in mammalian nuclei with enormous genomes and nucleosomes that would seemlingly obstruct 1D sliding remains poorly understood. One problem with applying the 1D+3D view in mammalian cells is that the number of off-target sequences and non-specific DNA binding sites is orders of magnitude greater in mammals than in prokaryotes (Iwahara and Clore, 2006). The much higher ratio of non-specific to specific binding sites in mammalian nuclei compared to prokaryotes suggests that mammalian cells may have evolved distinct mechanisms for guiding DNA-binding proteins towards their specific, cognate target sites. Indeed, recent work has suggested that non-specific binding to DNA can be a strong determinant of the distribution and nuclear exploration of DNA-binding proteins (McSwiggen et al., 2018).

Here we investigate how mammalian DNA-binding proteins find their nuclear targets and focus on the nuclear target-search mechanism of CTCF. CTCF, together with the cohesin complex, folds mammalian genomes into spatial domains known as Topologically Associating Domains (TADs), which regulate enhancer-promoter contacts and gene expression (Hansen et al., 2018a; Merkenschlager and Nora, 2016; Rada-Iglesias et al., 2018; Rowley and Corces, 2018). CTCF is an 11-Zinc Finger DNA-binding protein with unstructured N- and C-terminal domains, whose function remains poorly understood (Ghirlando and Felsenfeld, 2016). Cohesin is thought to be a ring-shaped multi-subunit complex, which holds together CTCF-demarcated TADs as chromatin loops (Hassler et al., 2018). Although CTCF and cohesin have emerged as key regulators of genome organization and function, their nuclear target search mechanisms have not been studied.

Using Single-Particle Tracking (SPT) and theoretical modeling, here we show that both CTCF and cohesin exhibit unusual nuclear target search mechanisms, where anisotropic diffusion arises from being transiently trapped in specific zones with a characteristic size of ∼200 nm. Our results indicate that these zones correspond to CTCF clusters and, surprisingly, we find that trapping inside them is largely due to CTCF’s RNA-binding region (RBR). Functionally, transient RBR-mediated trapping in zones increases the efficiency of CTCF’s DNA-target search mechanism by about 2.5-fold. More generally, we suggest that RNA-mediated “guiding” could be an effective way to control and regulate the local concentration of proteins in the nucleus around specific sites.

## RESULTS

### CTCF exhibits highly anisotropic nuclear diffusion

Our initial imaging analysis revealed that CTCF exhibits highly anomalous diffusion inside the nucleus (Figure 1-Figure Supplement 1A; Video 1-4). Motivated by this observation, we decided to systematically investigate how CTCF, cohesin and other nuclear proteins explore the mammalian nucleus using single-particle tracking (SPT). Taking advantage of our established mouse embryonic stem cell (mESC) and human osteosarcoma (U2OS) cell lines where CTCF and the cohesin subunit Rad21 have been endogenously HaloTagged (Hansen et al., 2017), we used stroboscopic photo-activation SPT (spaSPT; Figure 1A; (Hansen et al., 2017, 2018b)) to track single protein molecules over time in live cells. spaSPT overcomes common biases in SPT by using stroboscopic excitation to minimize “motion-blur” bias and by using photo-activation to track at low densities of ∼0.5-1.0 molecules per nucleus per frame, which minimizes tracking errors (Elf et al., 2007; Grimm et al., 2016; Hansen et al., 2017, 2018b; Manley et al., 2008). Most nuclear proteins exist in either a “bound” chromatin-associated state (e.g. trajectory A in Figure 1B) or a seemingly “free” diffusing state (e.g. trajectory B in Figure 1B). To explore the nuclear diffusion mechanism, we need to analyze exclusively the diffusive trajectory segments. We achieved this using a Hidden-Markov Model (HMM) (Persson et al., 2013) to classify trajectories into bound and free segments and then removed the bound segments and calculated the angle (Burov et al., 2013; Izeddin et al., 2014; Liao et al., 2012) between 3 consecutive localizations provided that both displacements making up the angle were much longer than our localization error of ∼35 nm (Figure 1B; full details in Materials and Methods). Finally, to comprehensively analyze the data at multiple spatiotemporal scales, we generated a very large data set by imaging 1,693 single cells at three different frame-rates (223 Hz, 134 Hz, 74 Hz; see Data Availability for raw data).

**Figure 1.**
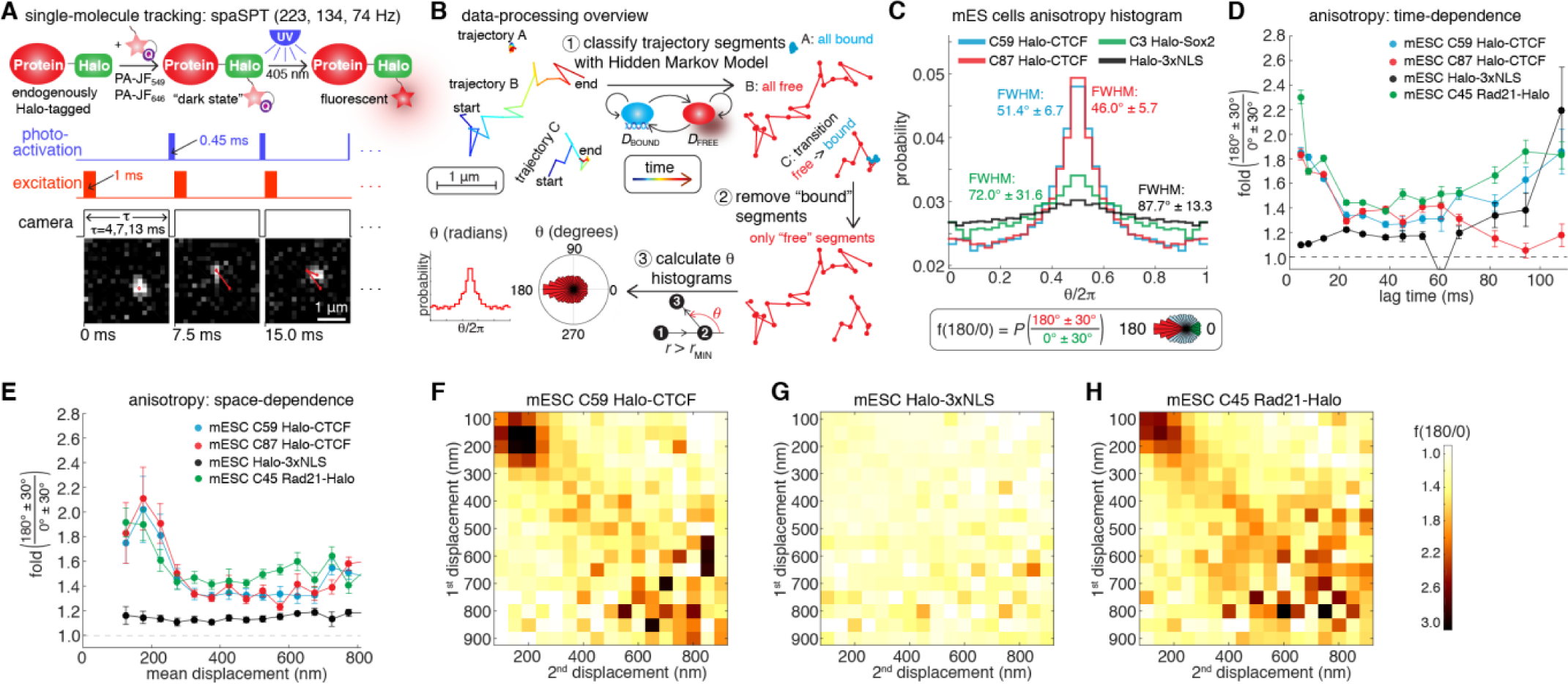
spaSPT reveals anisotropic CTCF diffusion in the nucleus. (**A**) Overview of spaSPT. Sketch of Halo-labeling with PA-JF_549_/PA-JF_646_, photo-activation, and overview of laser pattern. Below, representative raw images with tracking. (**B**) Overview of data-processing. spaSPT trajectories are classified into “bound” and “free” segments using an HMM and angles are only calculated from “free” segments where both displacements are above 200 nm. (**C**) Bulk histograms of the distribution of angles for CTCF (C59, C87), Sox2 (C3) and Halo-3xNLS in mESCs. Definition of fold-anisotropy, *f*_180/0_. (**D**) Plot of *f*_180/0_ vs. lag time averaging over all displacement lengths. (**E**) Plot of *f*_180/0_ vs. mean displacement length averaging over all lag times. (F-H) Anisotropy heatmaps showing *f*_180/0_ vs. length of the first and second displacement for C59 Halo-CTCF (**F**), Halo-3xNLS (**G**), C45 mRad21-Halo (**H**; in S/G2 phase of the cell cycle), averaging over all lag times. **Figure 1-Figure Supplement 1.** MSD-analysis and control Brownian motion simulations. **Figure 1-Figure Supplement 2.** 2-color spaSPT experiments to estimate the misconnection probability during tracking. **Figure 1-Figure Supplement 3.** Anisotropy for CTCF in human U2OS cells and cohesin in mESCs.

We began by analyzing two independent mESC Halo-CTCF clones, C59 and C87. The distribution of angles from diffusing CTCF trajectories showed a large peak at ∼180? (Figure 1C). This is surprising because it indicates that CTCF displays a directional bias: once CTCF has moved in one direction, it is significantly more likely to move backward in the opposite direction than it is to continue forward. This anisotropic diffusion behavior is not what one would expect for Brownian motion, which should have no directional bias. To quantify this effect, we use the fold anisotropy, *f*_180/0_: how many fold more likely is a step backward relative to a step forward, which for CTCF was 1.77 in mESCs. While confinement inside the nucleus is expected to cause some anisotropy due to e.g. collisions with the nuclear envelope, the free Halo-3xNLS protein was almost isotropic, *f*_180/0_=1.12, ruling out this as an explanation (Figure 1C). Could this be a general effect of all DNA-binding proteins? To test this, we analyzed a Halo-Sox2 knock-in mESC line (Teves et al., 2016), but Sox2 was also much less anisotropic than CTCF (*f*_180/0_=1.27; Figure 1C). Thus, CTCF exhibits unusual anisotropic diffusion behavior.

A previous study found that PTEFb exhibits scale-free anisotropic diffusion, with a magnitude of anisotropy that remains constant in space and time (Izeddin et al., 2014). To see if a similar mechanism holds for CTCF, we analyzed *f*_180/0_ as a function of the lag time between frames (Figure 1D) and the mean displacement length (Figure 1E). After an intial decline, *f*_180/0_ was relatively constant in time and somewhat noisy at long time-lags (Figure 1D). In contrast, CTCF showed a clear anisotropy peak at ∼200 nm displacements (Figure 1E). This was surprising since this type of diffusion has, to the best of our knowledge, not previously been observed. The spatial dependence was even clearer when we plotted *f*_180/0_ as a function of the length of the first and second displacements: CTCF showed a prominent peak (Figure 1F), which was not seen for Halo-3xNLS (Figure 1G). Using simulations, we verified that the observed anisotropy was not due to our localization uncertainty of ∼35 nm (Figure 1-Figure Supplement 1B-D). To further confirm that this result was also not due to tracking errors, we performed 2-color spaSPT control experiments: we labeled Halo-CTCF 1:1 with two distinguishable dyes, PA-JF_549_ and PA-JF_646_, which enabled us to identify any tracking errors (e.g. red→green switches within the same trajectory) and analyzed how tracking errors depend on density and displacement lengths. As expected, tracking errors were almost non-existent at the low densities of our imaging (Figure 1-Figure Supplement 2). However, we found that tracking errors increase *exponentially* with displacement length reaching ∼5% around ∼800 nm displacements (Figure 1-Figure Supplement 2A,F), and we thus limited our analysis to displacements below this length (full discussion in Appendix 1). Finally, we asked whether this mechanism is conserved between species and analyzed CTCF diffusion in human U2OS cells. Like in mESCs, human CTCF showed highly anisotropic diffusion with a large peak at ∼200 nm displacements (Figure 1-Figure Supplement 3A-D). Notably, although Sox2 did not, when we analyzed cohesin anisotropy, we found a similar anisotropic diffusion mechanism with a peak around ∼200 nm displacements (Figure 1D,E,H and Figure 1-Figure Supplement 3E-J). In summary, we conclude that CTCF and cohesin exhibit a highly unusual mechanism of anisotropic nuclear diffusion with a clear spatial dependence that is specific and conserved between mouse stem cells and differentiated human cells.

### A model featuring transient trapping in domains/zones can explain CTCF’s behavior

We next sought to understand the underlying mechanism of anisotropic diffusion by CTCF. A previous study explained anisotropic diffusion of the transcription complex PTEFb by suggesting that PTEFb moves along a fractal structure, most likely chromatin in the nucleus, and when it reaches a dead-end, must turn back (Izeddin et al., 2014). Since CTCF anisotropy is not uniform with displacement length (*i.e.,* it peaks at ∼200 nm), a similar fractal model with scale-free behavior cannot explain CTCF’s diffusion. We thus instead hypothesized that weak and transient interactions likely govern CTCF motion in the nucleus. If the probability of binding transiently at a given position was uniform throughout the nucleus, the resulting protein diffusion should be isotropic and Brownian, but with an effective (reduced) diffusion coefficient. However, if the source of CTCF anisotropic diffusion is transient localized trapping, such retention should occur in a domain/zone of a characteristic size (Figure 2A).

**Figure 2.**
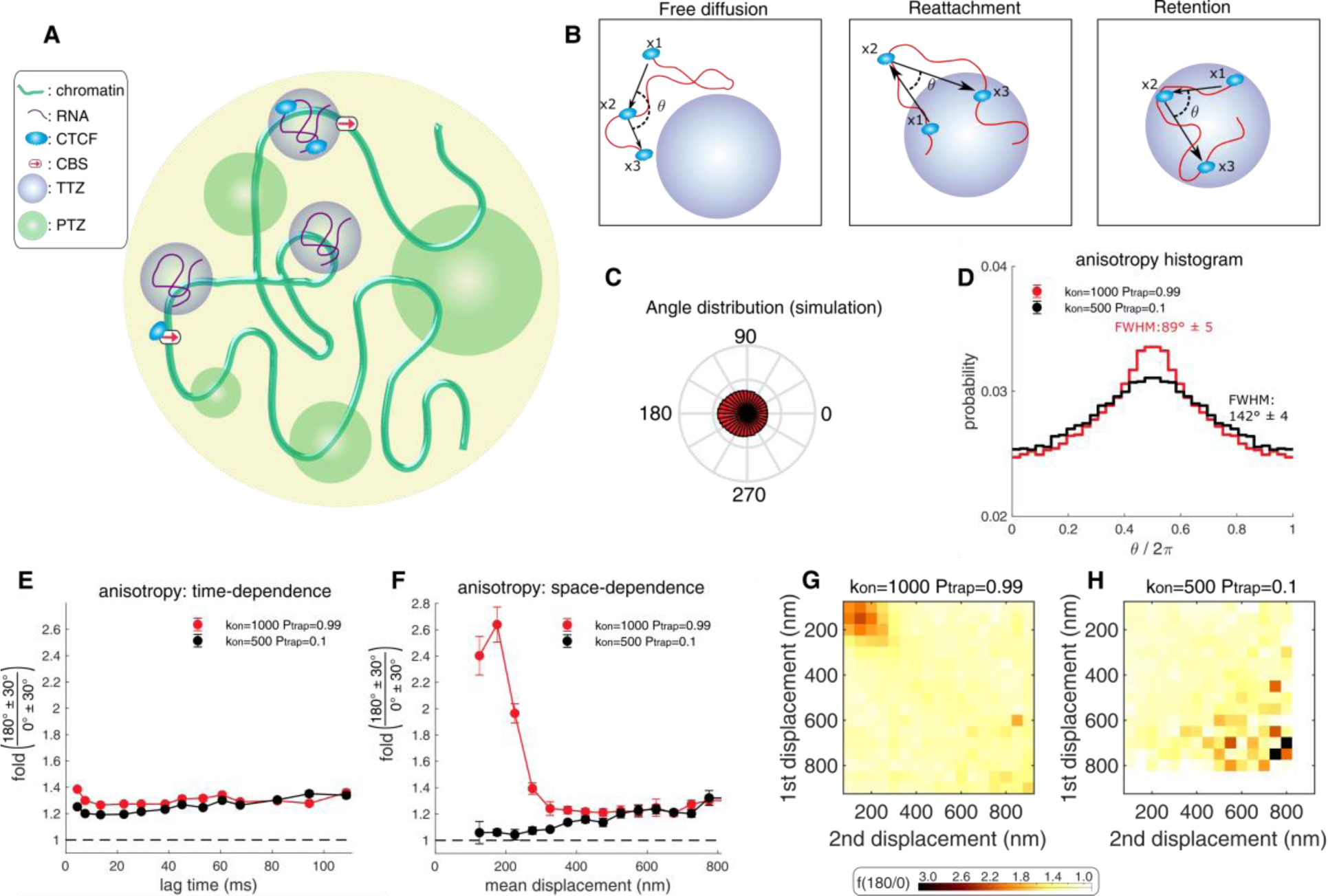
A model wherein CTCF diffusion in the nucleus is governed by its interaction with binding zones can explain the experimental data. (**A**) CTCF (light blue ellipse) diffuses inside the nucleus. It interacts with one of three zone types: (1) small, Cognate DNA Binding Sites (CBSs) (red arrow). (2) Transiently trapping zones (TTZs) of size 200nm (purple). (3) Trapping zones of different dimensions, whose size is derived from a power law distribution (Power-law distributed Trapping Zones – PTZs – green; for details in PTZs, please see Figure 3 and below). (**B**) The protein (light blue) is observed at 3 time points (x1, x2, x3) along its stochastic trajectory (red curve). Left: Outside of the zones, CTCF performs free diffusion. The angle of two consecutive steps is uniformly distributed for this mode of motion. Middle: When CTCF leaves a TTZ (x1), diffuses freely (x2), and then reattaches back to the TTZ (x3), the angle distribution is anisotropic. Right: While inside a zone, CTCF can be reflected from its boundary. The angle distribution, in this case, will be anisotropic (**C**) The angle distribution for a protein performing diffusion in a nucleus containing only CBSs and TTZs. See Table S1 for the simulation parameters. (**D**) Angle distribution of the trajectory computed from the simulation when the nucleus contains only zones of type 1 and 2 (CBSs and TTZs). CTCF is trapped with high probability (*P*_*trap*_ = 0.99) when touching the zones (red curve). While inside the zone, it can transiently arrest its motion with a high rate *k*_*on*_ (black curve). The protein can be trapped with low probability (*P*_*trap*_ = 0.1) when touching the zone. While inside the zone, it can transiently arrest its motion with smaller rate. (**E**) Plot of *f*_180/0_ vs. lag time averaging over all displacement lengths of the simulation data shown in **D**. (**F**) Plot of *f*_180/0_ vs. mean displacement length averaging over all lag times. (**G-H**) Anisotropy heatmaps showing *f*_180/0_ vs. length of the first and second displacement for high binding probability (*P*_*trap*_ = 0.99, *k*_*on*_ = 1000) (**G**) and for smaller binding probability (*P*_*trap*_ = 0.1, *k*_*on*_ = 500) (**H**). **Figure 2-Figure Supplement 1.** Overview of the model. **Figure 2-Figure Supplement 2.** MSD from model simulations.

We tested this hypothesis by simulating chromatin as self-avoiding polymers confined inside the nucleus (Amitai et al., 2017) (Figure 2-Figure Supplement 1A) and simulated protein motion and interaction with chromatin (see (Amitai, 2018) and Materials and Methods for full details). The protein undergoes Brownian motion inside the nucleus but upon encountering a Transiently Trapping Zone (monomer; TTZ) has a certain probability *P*_*trap*_ of becoming absorbed and trapped. We model TTZs as spherical. These zones trap CTCF transiently with a time-scale comparable to our experimental frame rates and thus result in trajectories that appear as both free and bound, depending on the bound state of the protein. CTCF also binds strongly to Cognate DNA Binding Sites (CBSs) (Figure 2A) with a residence time for specific binding of ∼1-4 minutes (Hansen et al., 2017), which is effectively infinitely stable on the time-scale of our SPT experiments (milliseconds).

We thus considered three kinds of interactions. The first class, CBSs, accounts for stable binding of CTCF. The protein remains bound there for a characteristic time *τ*_*CBS*_ = 1 min. The second class, TTZs, traps the protein for a much shorter characteristic time *τ*_*TB*_ (*τ*_*CBS*_ ≫ *τ*_*TTZ*_) (Figure 2A). While trapped, the particle is free to diffuse inside the spherical TTZs that have a radius *ε*_*TTZ*_. When the diffusing protein hits the boundary, it can exit the TTZ with probability *P*_*exit*_ or be reflected. Upon exit, the particle is placed at a distance *a*_*TTZ*_ from the TTZ center and starts diffusing again (Figure 2-Figure Supplement 4B). The model parameters are chosen such that the protein has a probability to escape a zone upon release rather than rebinding immediately. Upon encountering another TTZ (arriving at a distance *ε*_TTZ_ from its center), it has a probability to yet again become trapped. While diffusing inside the TTZ, the protein can transiently bind at any position with Poissonian on-rate *k*_on_ and then be released with Poissonian off-rate *k*_*off*_. We will discuss a third class of zones, Power-law distributed Trapping Zones (PTZs), is further details below.

We then tested if our model could reproduce our experimental observations. We simulated Brownian diffusion using Euler’s scheme (Schuss, 2009) subject to cognate and transient interactions (full details in Materials and Methods). We chose the simulation parameters including diffusion coefficient, total bound fraction and localization error to approximately match our experimental measurements. In the experiments, a CTCF molecule stably bound to a CBS appears to be immobile throughout its trajectory and we therefore simulate CBS binding as immobile subject to 35 nm localization uncertainty. We note that stably bound CTCF will thus be filtered out by our analysis pipeline (Figure 1B), but we nevertheless simulate stably bound CTCF to match the experimental data. We take the size of the TTZs to be *ε*_TTZ_ = 200nm. Although protein diffusion is Brownian in our model, transient trapping of CTCF in the TTZs faithfully reproduced our experimental observations, including high anisotropy (Figure 2C-D) and a clear peak of anisotropy at ∼200 nm mean displacements (Figure 2F) and a relatively constant trend in time (Figure 1D; Figure 1-Figure Supplement 3B). The relatively constant value of *f*_180/0_ in time (Figure 2E) suggests that the interaction time of a protein with a zone has a broad distribution. Interestingly, we note that the location of the peak in Figure 2G provides information about the size of the TTZ. Our simulations reproduced this behavior at high trapping probability – if we lowered the probability *P*_*trap*_ of trapping CTCF in these zones, the diffusion remained anisotropic, but the anisotropy peak largely disappeared (Figure 2F). While we considered a large variety of models and parameter combinations, only models with zones of a fairly well-defined size that trap CTCF with high probability could reproduce the experimental data. We refer to this model as Anisotropic Diffusion through transient Trapping in Zones (ADTZ).

What is the underlying mechanism of ADTZ? When a protein transiently interacts with chromatin, its dynamics will be governed by rapid reattachment to the release site (Amitai, 2018). Indeed, if a protein is bound to a site for a given time, upon disassociation it is much more likely to re-attach to the same site rather than bind to another site. Protein diffusion therefore is biased toward its starting position. When the characteristic time to return is on the order of the frame-rate of our SPT experiments (milliseconds) the protein will appear to take a step back following a forward step. We call this mechanism that will contribute to the angle anisotropy *reattachment* (Figure 2B). While the protein is trapped in the TTZ it is reflected from the domain boundary with some probability. We call this mechanism *retention* (Figure 2B). Hence, we suggest that the origin of anisotropy could be two-fold and originate from a combination of *reattachment* and *retention*.

Finally, diffusion is often analyzed using mean square displacement (MSD) analysis. Many nuclear proteins exhibit subdiffusive behaviors (Metzler et al., 2014; Saxton, 2007), where their MSD grows as a power law with time *MSD*_*i*_ (*τ*) = (*r*_*i*_ (*t* + *τ*) – *r*_*i*_ (*t*))*∼τ^α^*, with *α* < 1. By analyzing the trajectories, we found that CTCF exhibits anomalous diffusion with an exponent of *α∼*0.8 (Figure 1-Figure Supplement 1A; value is sensitive to conditions). The observed anomalous diffusion has been attributed to motion in a crowded environment (Höfling and Franosch, 2013) or diffusion in a hypothetical fractal domain of the nucleus (Bancaud et al., 2009). This motion is often phenomenologically modeled as a continuous time random walk (CTRW), where a particle is trapped for a time drawn from a long tailed power-law distribution (Metzler and Klafter, 2000). To examine the consequences of the ADTZ model on the dynamics of a protein, we computed the MSD of our simulation. We find that the protein appears to perform subdiffusive motion with an anomalous exponent of about *α∼*0.89 (Figure 2-Figure Supplement 2; value is sensitive to conditions) due to the rapid reattachment at the release site. The value of *α* is not universal and depends on model parameters. Interestingly, higher reattachment probability leads to a lower anomalous exponent. A freely diffusing particle confined inside the nucleus has an anomalous exponent *α* = 0.97, which is smaller than 1 because of the nuclear confinement. Thus, the ADTZ model can mechanistically explain the phenomenology of subdiffusive protein motion in the nucleus without invoking power-law dissociation times as is assumed in CTRW (Metzler and Klafter, 2000).

### Distinct interactions by CTCF’s RBR- and ZF-domains control CTCF diffusion

Our simulations suggest an intriguing mechanistic model (ADTZ) where zones of an unknown nature transiently trap diffusing CTCF with high probability. What could be the nature of these TTZs? We previously demonstrated using super-resolution PALM imaging that CTCF and cohesin form small co-localizing clusters in the nucleus (Hansen et al., 2017). Here, we hypothesize that the TTZs could correspond to CTCF clusters, since clustering is due to self-interaction and self-interaction might also explain transient CTCF trapping. In our companion paper (Hansen et al., 2018c), we show that CTCF self-association is largely RNA-mediated and DNA-independent consistent with a previous report (Saldaña-Meyer et al., 2014) and that CTCF clustering is substantially reduced after endogenous deletion of CTCF RNA-binding region (RBR; mESC C59D2 ΔRBR-Halo-CTCF). Thus, if the ADTZ model hypothesis is correct and TTZs correspond to CTCF clusters, it should be possible to abolish TTZ-mediated trapping of CTCF in two ways: a) by reducing the trapping probability of CTCF (Figure 2F) or b) by reducing the number of CTCF clusters.

To test these hypotheses and elucidate the molecular mechanism of CTCF diffusion we performed spaSPT for a series of CTCF mutants in mESCs (Figure 3A-F) and U2OS cells (Figure 3-Figure Supplement 1). Whereas a full deletion of CTCF 11 Zinc Finger domain (ΔZF), which is required for DNA-binding (Hansen et al., 2017; Nakahashi et al., 2013), caused CTCF diffusion to become nearly isotropic, ΔRBR-CTCF remained almost as anisotropic as wt-CTCF at the bulk level (Figure 3A). However, analysis of the fold–anisotropy (*f*_180/0_) as a function of time (Figure 3B) and space (Figure 3C) revealed two surprising results. First, the anisotropy peak largely, though not entirely, disappeared in ΔRBR-CTCF (Figure 3C-E). Our ADTZ theory (Figure 2) predicts that this can occur in two ways: a) by reducing the number of TTZs (Figure 3-Figure Supplement 2) and b) by reducing the trapping probability (Figure 2A-B), to the TTZs. Clustering (TTZs) is clearly reduced in ΔRBR-CTCF (Hansen et al., 2018c) and it is likely that *P*_*trap*_ is too. These results paint a mechanistic picture for the ADTZ model: the RBR domain mediates CTCF clustering, which serve as TTZs that transiently trap diffusing CTCF in zones of a defined size (∼200 nm), resulting in highly anisotropic CTCF diffusion. However, this points to the second surprising result: ΔRBR-CTCF diffusion is still highly anisotropic (Figure 3A), but with a magnitude (*f*_180/0_) that is approximately constant in both time and space (Figure 3B-C). Thus, without the RBR, CTCF exhibits approximately scale-free anisotropy similar to PTEFb (Izeddin et al., 2014).

**Figure 3.**
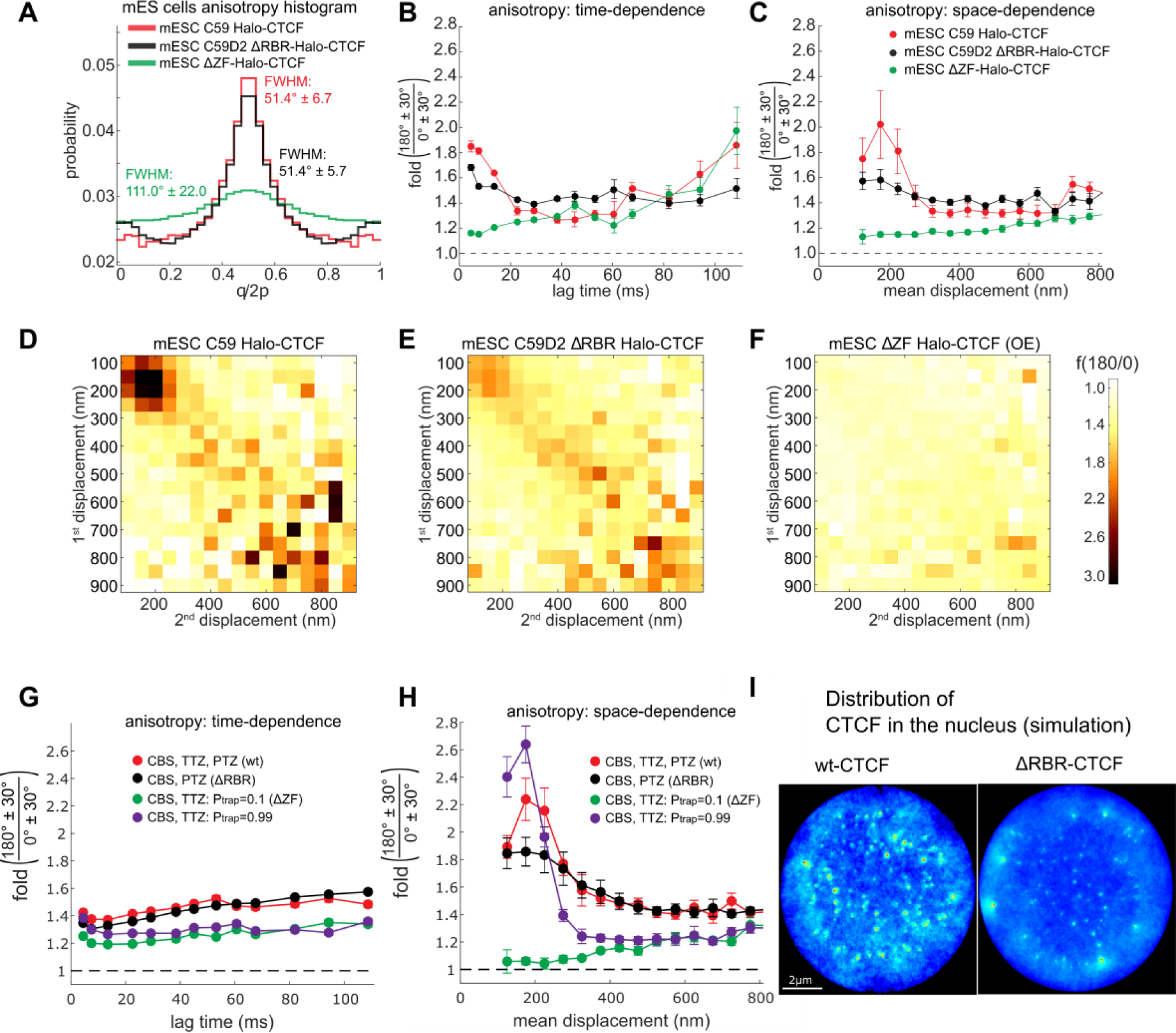
Anisotropy and nuclear distribution of Δ RBR-CTCF. (**A**) Bulk angle distribution histograms for CTCF (C59) (red), CTCF mutant that does not have the RBR domain and does not bind RNA (C59D2) (black), CTCF with a deletion of CTCF’s 11 Zinc Finger domain (ΔZF) and which does not bind both DNA and RNA (ΔZF-Halo-CTCF) (green). (**B**) Plot of *f*_180/0_ vs. lag time averaging over all displacement lengths. (**C**) Plot of *f*_180/0_ vs. mean displacement length averaging over all lag times. (**D-F**) Anisotropy heatmaps showing *f*_180/0_ vs. length of the first and second displacement for the three cell lines, averaging over all lag times. **Simulations.** (**G-H**) Plot of *f*_180/0_ vs. lag time averaging over all displacement lengths (**G**) or vs. mean displacement length averaging over all lag times (**H**). The full model representing the wt, where CTCF interacts with high trapping probability with all three zones (CBS, TTZ, PTZ – see Figure 2A) (red). CTCF interacts with high trapping probability with CBS and PTZ but has a very weak affinity for the TTZ (black) (representing ΔRBR-CTCF). CTCF has a very low affinity to the CBSs and TTZs. There are no PTZs (green) (representing ΔZF-CTCF). CTCF has high affinity towards CBSs and TTZs. There are no PTZs (purple). (**I**) The distribution of CTCF in the nucleus estimated from the simulation (full model corresponding to wt-CTCF) (left), and the model where CTCF weakly interacts with TTZ (corresponding to ΔRBR-CTCF) (right). **Figure 3-Figure Supplement 1.** Anisotropy wild-type and mutant CTCF in human U2OS cells. **Figure 3-Figure Supplement 2.** Effect of TTZs abundance on anisotropy.

Since ΔZF-CTCF is essentially isotropic, these results suggest a more complicated model wherein CTCF anisotropy arises through a combination of 2 mechanisms. First, RBR mediated interactions transiently trap CTCF in small TTZs, resulting in a peak of anisotropy at ∼200 nm. Second, Zinc Finger (ZF) mediated interactions, perhaps due to transient DNA-interactions, generally trap CTCF without a clearly defined scale. Lending further support to this interpretation, we observed the same ΔZF-and ΔRBR-mutant and wild-type CTCF behavior in human U2OS cells (Figure 3-Figure Supplement 1).

Since our model with TTZs already reproduces anisotropy at a defined scale, we next studied whether adding additional zones could explain the scale-free anisotropy observed without the RBR (Figure 3A-E). Because the data suggests a “scale-free” anisotropy, which is due to transient DNA-interactions, we added to the nucleus zones of different sizes (Figure 2A) which are sampled from a power law distribution (Figure 3H) (*P*_*PTZ*_ (*ε*)*∼ε*^-γ^) and term theses zones *Power-law Trapping Zones* (PTZs). To reproduce our experiments, we choose the PTZs such that large zones are rarer and trap the protein less efficiently (specifically *P*_*trap,PTZ*_(*ε*)*∼ε*^-*δ*^; see Materials and Methods). For very small zones, the theory suggests that for *δ* of order 1, we would get scale-free anisotropy. We then performed Brownian simulations of an interacting protein where chromatin was represented by a polymer where the majority of its monomers are TTZs (of size 200nm), a smaller fraction is PTZs (with a Power Law size distribution), and a third fraction consists of CBSs. The results recapitulate both the peak of anisotropy at ∼200 nm and the large (*f*_180/0_∼1.5) and scale-free anisotropy at longer displacements (Figure 3G-H). Thus, the ADTZ model can explain the behavior of wt-CTCF and when we “computationally mutate” the ADTZ-part, the model can also explain the behavior of ΔRBR-CTCF. Finally, we computationally estimated the distribution and clustering of wt-CTCF and ΔRBR-CTCF in the nucleus as it results from our simulations (Figure 3I). Notably, the computationally estimated difference in distribution between wt-CTCF and ΔRBR-CTCF looks qualitatively similar to the distribution observed in super-resolution PALM experiments (Hansen et al., 2018c).

### CTCF target search mechanism is RBR-guided

Having elucidated a surprisingly complicated mode of CTCF diffusion, we next asked what the function might be. We reasoned that cells might regulate the diffusion of DNA-binding proteins in order to regulate the efficiency of their cognate target site search. To study CTCF target-search efficiency, we performed spaSPT and Fluorescence Recovery After Photobleaching (FRAP) experiments. First, spaSPT experiments coupled with 2-state model-based analysis (Hansen et al., 2017, 2018b) revealed that the free diffusion coefficient (*D*_FREE_) was apparently unaffected, whereas the total bound fraction decreased significantly from ∼62.4% to 42.0% (Figure 4A-C). Since our spaSPT experiments capture both non-specifically “bound” CTCF and specifically bound CTCF (characterized by a ∼1-4 min residence time (Hansen et al., 2017)), we next asked using Fluorescence Recovery After Photobleaching (FRAP) experiments which population was affected. FRAP independently confirmed that the bound fraction was substantially reduced (Figure 4D), but model fitting revealed that the apparent residence time of specifically bound CTCF was not significantly affected, though the population size of specifically bound CTCF was reduced (Figure 4-Figure Supplement 1). Thus, these results demonstrate a surprising function of CTCF RBR: the strength of CTCF specific binding to cognate sites is unaffected by the RBR, but the amount of CTCF engaged in specific binding is strongly impacted. In a simplified 2-state model, the fraction of CTCF specifically bound to chromatin is controlled by the balance between its binding (ON) and dissociation (OFF) rates 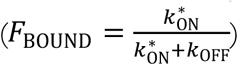. Within this framework and using our measurements, we find that the RBR increases 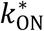 by ∼2.5-fold (see Materials and Methods for calculation). The ON-rate is the inverse of the time it takes the protein to find its cognate binding site (search time). Thus, the RBR increases the frequency or rate of CTCF finding a specific DNA-binding site by ∼2.5-fold, without affecting CTCF affinity for specific binding sites. This suggests that the ADTZ-diffusion mechanism serves to increase the efficiency of CTCF target search mechanism. Additionally, this suggests that TTZs serve to “load” or “guide” CTCF towards cognate DNA binding sites. Hence, CTCF cognate binding sites may be located within or near TTZs. If this were the case, mechanistically, the TTZs would increase the local concentration of CTCF around its cognate DNA binding sites, thus increasing the local ON-rate.

**Figure 4.**
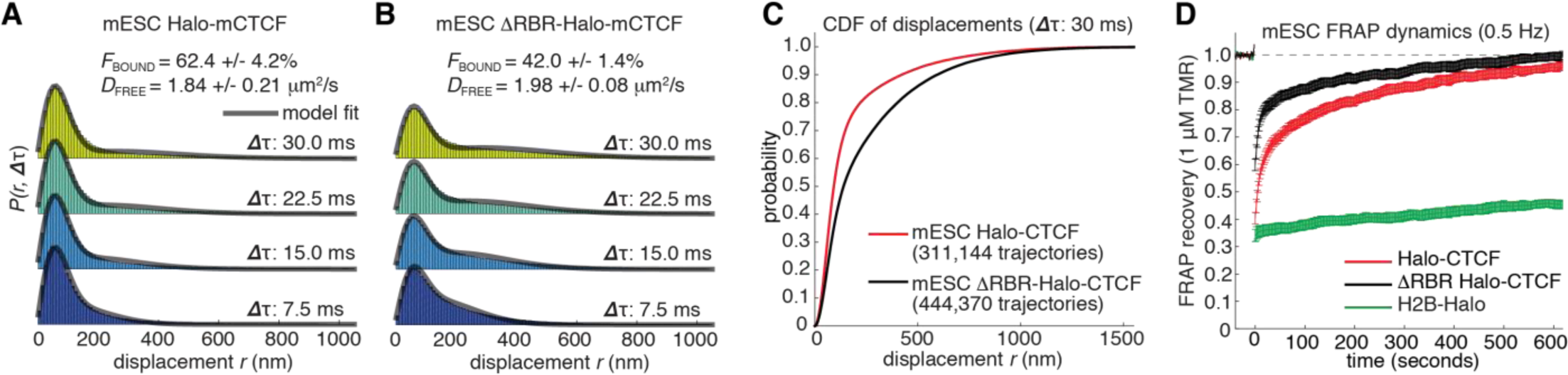
RBR-guided CTCF target search mechanism. **(A-B**): spaSPT histograms of displacements for wt-CTCF (**A**) and ΔRBR-CTCF (**B**). Raw displacement data for four different lag times are shown with a 2-state Spot-On model fit (bound vs. free) overlaid (Hansen et al., 2017, 2018b). The inferred bound fraction and free diffusion coefficients are shown. Standard errors (+/-) are indicated. (**C**) Cumulative probability function of displacement lengths (Δ*τ*=30 ms) for the same data as in **A-B**. (**D**) Fluorescence Recovery After Photobleaching (FRAP) data for wt-CTCF, ΔRBR-CTCF, and histone H2B (control). Model-fits and inferred residence times are shown in Figure 4-Figure Supplement 1. **Figure 4-Figure Supplement 1.** Model-fit to FRAP data.

## DISCUSSION

### Anisotropic diffusion through transient and weak interactions in the nucleus

It has long been known that diffusion in cells can be anomalous or non-Brownian (Metzler et al., 2014) and anomalous sub-diffusive behavior has previously been reported for DNA (Bronstein et al., 2009), RNA (Golding and Cox, 2006), and proteins (Izeddin et al., 2014). Here, we introduce two new concepts and a new approach for studying the diffusion of nuclear proteins in live cells. Technically, by largely eliminating tracking errors and motion-blur artifacts, spaSPT (Hansen et al., 2017, 2018b) overcomes common technical artifacts in SPT that can bias the analysis, and our analysis pipeline (Figure 1B) crucially filters out chromatin-associated trajectories. While we have focused here on analyzing the angle anisotropy, we anticipate that our analysis has only scratched the tip of the iceberg and we suggest that our approach can usefully be applied to many other nuclear proteins.

Conceptually, first, our work has revealed a new mode of nuclear exploration, which results in anisotropic motion: the Anisotropic Diffusion through transient Trapping in Zones (ADTZ) model (Figure 2). Unlike phenomenological models of sub-diffusion such as CTRW (Metzler et al., 2014), which assumes sub-diffusion is the result of power-law waiting time distributions at a binding site, our model offers a geometrical and mechanistic explanation for CTCF dynamics. Additionally, CTRW is not expected to result in anisotropic motion. We suggest that the primary contribution to wt-CTCF anisotropic dynamics is its interactions within RBR-mediated TTZs of a characteristic ∼200-nm size (Figure 5A). Within a zone, CTCF is allowed to move and participate in transient interactions, which effectively reduce its in-zone diffusion coefficient. The ADTZ model can mechanistically explain sub-diffusion as coming from the retention and reattachment mechanisms (Figure 2). While inside the zone, CTCF is reflected from its boundary with a certain probability (*retention*), resulting in a characteristic residence time within the zone. Once CTCF escapes one zone, it is very likely to rebind to the same zone because of its proximity (*reattachment*), which results in anisotropic behavior. Thus, the ADTZ model suggests that CTCF anisotropy results from the non-uniform distribution of zones inside the nucleus. Mechanistically, our results suggest that the transient trapping zones (TTZs) likely correspond to clusters of CTCF and that the trapping is mediated by CTCF RBR domain. Speculatively, we suggest that if RNA(s) hold together CTCF clusters, weak CTCF-RNA interactions mediated by the RBR-domain may repeatedly trap CTCF. Identifying such putative RNA(s) will be an important future direction. It will be interesting to explore in the future if other proteins exhibit similar mechanisms – especially for proteins that also form clusters or hubs (Mir et al., 2018).

**Figure 5.**
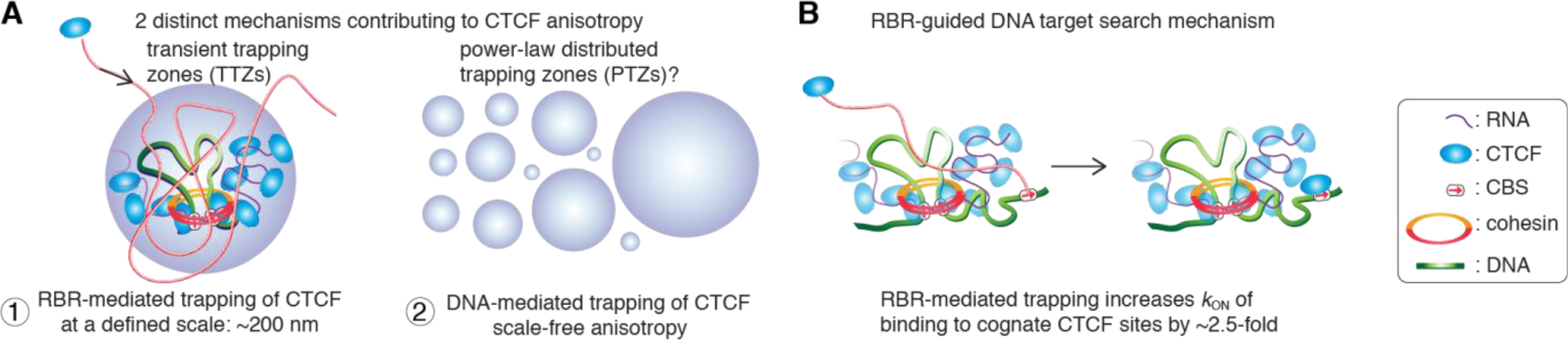
Model. (**A**) Two distinct mechanisms contribute to CTCF anisotropy. First, RBR-mediated trapping of CTCF in transiently trapping zones (TTZs) of a characteristic size (∼200 nm). TTZs are likely to correspond to clusters of CTCF that form in a largely RBR-dependent manner and therefore, presumably, are held together by RNA(s). Cognate Binding Sites (CBSs) reside within the TTZ, as does a piece of DNA strand (green). Second, ΔRBR-CTCF still displays scale-free anisotropy, which is presumably due to non-specific interactions with DNA. We speculatively model this as arising from trapping in power-law distributed zones (PTZs). (**B**) Target-search mechanism. The RBR increases the apparent rate at which CTCF locates a cognate DNA-binding site. Given that the RBR also mediates CTCF clustering, we speculate that RNA-mediated CTCF clusters, perhaps near loop boundaries, help “guide” diffusing CTCF towards its cognate DNA binding sites (CBSs; DNA is shown in green).

Second, ΔRBR-CTCF still displays anisotropic diffusion with a scale-free magnitude, but CTCF without its Zinc Finger domains exhibit isotropic diffusion (Figure 3). Conceptually, this suggests that protein diffusion is modular, tunable and programmable (Woringer and Darzacq, 2018). Specifically, some protein domains exhibit essentially isotropic diffusion (e.g. the N- and C-terminal domains of CTCF, Halo-3xNLS etc.). CTCF’s RBR-domain mediates anisotropic diffusion through an ADTZ-type mechanism and CTCF’s 11-Zinc Finger-domain mediates anisotropic diffusion with a scale-free magnitude (Figure 3). Since ΔRBR-CTCF does not participate in transient interactions within the 200-nm zones, we suggest that the scale-free anisotropy observed for ΔRBR-CTCF is due to DNA-mediated interactions. This is because this anisotropy is lost when CTCF’s DNA-binding domain is lost (Figure 3). We model scale-free DNA-mediated anisotropy as originating from interaction with zones of different sizes, where the larger zones are less sticky (Figure 5A). In these zones (PTZs), which we speculate have a power-law distribution, CTCF experiences transient interactions as in the TTZs. While the origin of the PTZs is unknown at this point, these might correspond to TAD or compartment structures, which vary both in range and size. Moreover, we note that diffusion in a disordered medium could also explain scale-free anisotropy (Havlin and Ben-Avraham, 2002).

We conclude that both RNA- and DNA-mediated interactions govern CTCF diffusion in a modular manner and these interactions result in anisotropy that manifests itself at different scales. Thus, by mixing and matching protein domains with defined diffusion mechanisms, it should in principle be possible to design a protein with a desired diffusion mechanism. This could both be exploited in synthetic biology approaches to engineer proteins with desired target search mechanisms or by the cell during evolution to fine-tune function.

### Functional implications of RBR-mediated interactions

Although still speculative at this point, our results suggest a strong link between CTCF clustering, diffusion and target search mechanism (Figure 5B). Specifically, our results suggest that CTCF clusters correspond to TTZs. Since clusters largely form in an RBR-dependent manner (Hansen et al., 2018c), they are presumably held together by one or more species of RNA, as evidenced by co-IP experiments in our companion paper (Hansen et al., 2018c). Moreover, since the RBR increases the rate at which CTCF locates cognate DNA-binding sites by ∼2.5-fold, this suggests a model where the CTCF DNA-target search is guided by RNA and where CTCF clusters “guide” diffusing CTCF proteins towards nearby cognate DNA binding sites (Figure 5B). This model significantly changes the view on the mechanism of target search and protein localization in mammalian cells. Rather than the facilitated diffusion picture (alternating between 1D diffusion - e.g. by sliding along the DNA - and 3D diffusion), our model and experimental results suggest that protein localization and target search depend on the formation of local high concentration zones or clusters, perhaps around cognate binding sites, mediated by RNA. It will be interesting to explore if other proteins exhibit similar “guided” target search mechanisms.

CTCF and cohesin regulate genome organization together (Rowley and Corces, 2018). It is therefore striking that both CTCF (Figure 1) and cohesin (Figure 1H and Figure 1-Figure Supplement 3) exhibit similar ADTZ-type anomalous diffusion. The similar diffusion mechanisms are especially interesting in light of the fact that how and where cohesin is loaded onto the genome remains poorly understood (Busslinger et al., 2017). It will be interesting to explore in the future if TTZs also contribute to topological loading of cohesin on chromatin. Consistent with an RBR-connection between CTCF, cohesin and genome organization, we show in a companion paper that a significant fraction of CTCF loops are lost in ΔRBR-CTCF mESCs (Hansen et al., 2018c). Thus, the same protein domain simultaneously regulates CTCF diffusion, clustering, target search mechanism, and function. Our results highlight the power of SPT, theory, and analysis, when coupled with genome-editing and mutations, as an approach to discover protein domains engaging in important interactions. It will be exciting to apply a similar approach to other nuclear proteins in the future.

### Materials and Methods; Supplementary Figures

Detailed methods and Supplementary Figures and Tables have been uploaded as a separate Supplementary Information file.

## Supporting information

## Acknowledgements

We thank Luke Lavis for generously providing JF dyes, Maxime Woringer for insightful discussions and help with simSPT, Astou Tangara and Ana Robles for microscope assembly and maintenance, Gina Dailey for help and assistance with cloning, and Dr. Kartoosh Heydari at the Li Ka Shing Facility for flow cytometry assistance. We thank Andrew Seeber, Khanh Dao Duc, David McSwiggen, and other members of the Tjian and Darzacq labs for comments on the manuscript. This work was performed in part at the CRL Molecular Imaging Center, supported by the Gordon and Betty Moore Foundation. ASH is a postdoctoral fellow of the Siebel Stem Cell Institute. This work was supported by NIH grants UO1-EB021236 and U54-DK107980 (XD), the California Institute of Regenerative Medicine grant LA1-08013 (XD), by the Howard Hughes Medical Institute (003061, RT), and Massachusetts General Hospital grant 214931 (AA). We thank Sheila S Teves for providing the preprint template.

