## Supplementary material for "Guided nuclear exploration increases CTCF target search efficiency"

#### MATERIALS AND METHODS

##### Cell culture

JM8.N4 mouse embryonic stem cells (Pettitt et al., 2009) (male mESCs; Research Resource Identifier: RRID:CVCL_J962; obtained from the KOMP Repository at UC Davis) and human U2OS osteosarcoma cells (Research Resource Identifier: RRID:CVCL_0042) were grown and handled as described previously(Hansen et al., 2017). Briefly, mES cells were grown on plates pre-coated with a 0.1% autoclaved gelatin solution (Sigma-Aldrich, St. Louis, MO, G9391) under feeder free conditions in knock-out DMEM with 15% FBS and LIF (full recipe: 500 mL knockout DMEM (ThermoFisher, Waltham, MA, #10829018), 6 mL MEM NEAA (ThermoFisher #11140050), 6 mL GlutaMax (ThermoFisher #35050061), 5 mL Penicillin-streptomycin (ThermoFisher #15140122), 4.6 μL 2-mercapoethanol (Sigma-Aldrich M3148), 90 mL fetal bovine serum (HyClone Logan, UT, FBS SH30910.03 lot #AXJ47554)) and LIF. mES cells were fed by replacing half the medium with fresh medium daily and passaged every two days by trypsinization. The homozygously endogenously Halo-tagged cell lines (e.g. C45 mRad21-Halo) and the cell lines stably over-expressing a transgene (e.g. H2B-Halo or Halo-3xNLS) have been described previously as well (Hansen et al., 2017). Likewise, female U2OS cells (Research Resource Identifier: RRID:CVCL_0042) were grown in low glucose DMEM with 10% FBS (full recipe: 500 mL DMEM (ThermoFisher #10567014), 50 mL fetal bovine serum (HyClone FBS SH30910.03 lot #AXJ47554) and 5 mL Penicillin-streptomycin (ThermoFisher #15140122)) and were passaged every 2-4 days before reaching confluency. The homozygously endogenously Halo-tagged cell lines (e.g. C32 Halo-hCTCF) and the cell lines stably over-expressing a transgene (e.g. H2B-Halo or Halo-3xNLS) have been described previously as well (Hansen et al., 2017).

Transient transfection experiments were also performed as previously described (Hansen et al., 2017). Briefly, cells were plated on plasma-cleaned 25 mm circular no 1.5H cover glasses (Marienfeld, Germany, High-Precision 0117650) either directly (U2OS) or MatriGel coated (mESCs; Fisher Scientific, Hampton, NH, #08-774-552 according to manufacturer's instructions just prior to cell plating). After overnight growth and ~18-36 hours before the imaging experiment, cells were then transfected with plasmid encoding the protein of interest using 250-1000 ng of plasmid per well in a 6-well plate using Lipofectamine 3000 (ThermoFisher #L3000008). All plasmids were identical except for the protein of interest and are as follows: the protein of interest (e.g. wt-Halo-mCTCF or wt-mRad21-Halo) was expressed from a L30 promoter (we previously found L30 to only somewhat change the behavior compared to endogenously tagged proteins, unlike CMV or EF1a promoters which strongly changed the behavior) and followed by an SV40 poly(A) signal. Downstream of the protein of interest, EGFP-3xNLS was expressed from a PGK promoter and followed by a bGH poly(A) signal. GFP-NLS facilitated finding transfected cells and outlining the nucleus. Moreover, the GFP signal was generally proportional to the expression level of the protein of interest and therefore made it possible to identify cells with a robust, but not too high, expression level. In summary, for transient transfection experiments, cells were plated two days before the imaging experiment and transfected one day before the imaging experiment.

For live-cell imaging, the medium was identical except DMEM without phenol red was used (ThermoFisher #31053028). Both mouse ES and human U2OS cells were grown in a Sanyo copper alloy IncuSafe humidified incubator (MCO-18AIC(UV)) at 37°C/5.5% $CO_2$.

Cell lines were pathogen tested (IMPACT II test for mESC C59; PCR-based mycoplasma assay for U2OS C32) and authenticated through STR profiling (U2OS) as described previously (Hansen et al., 2017). All cell lines will be provided upon request.

##### Live cell single-molecule imaging and analysis (spaSPT)

The indicated cell line was grown overnight on plasma-cleaned 25 mm circular no 1.5H cover glasses (Marienfeld, Germany, High-Precision 0117650) either directly (U2OS) or MatriGel coated (mESCs; Fisher Scientific, Hampton, NH, #08-774-552 according to manufacturer's instructions just prior to cell plating). After overnight growth, cells were labeled with 5-50 nM PA-$JF_{549}$ or PA-$JF_{646}$(Grimm et al., 2016) for ~15-30 min and washed twice (one wash: medium removed; PBS wash; replenished with fresh medium). At the end of the final wash, the medium was changed to phenol red-free medium keeping all other aspects of the medium the same. Single-molecule imaging was performed on a custom-built Nikon TI microscope (Nikon Instruments Inc., Melville, NY) equipped with a 100x/NA 1.49 oil-immersion TIRF objective (Nikon apochromat CFI Apo TIRF 100x Oil), EM-CCD camera (Andor, Concord, MA, iXon Ultra 897; frame-transfer mode; vertical shift speed: 0.9 μs; -70°C), a perfect focusing system to correct for axial drift and motorized laser illumination (Ti-TIRF, Nikon), which allows an incident angle adjustment to achieve highly inclined and laminated optical sheet illumination (Tokunaga et al., 2008). The incubation chamber maintained a humidified 37°C atmosphere with 5% $CO_2$ and the objective was also heated to 37°C. Excitation was achieved using the following laser lines: 561 nm (1 W, Genesis Coherent, Santa Clara, CA) for PA-$JF_{549}$; 633 nm (1 W, Genesis Coherent, Pala Alto, CA) for PA-$JF_{646}$; 405 nm (140 mW, OBIS, Coherent) for all photo-activation experiments. The excitation lasers were modulated by an acousto-optic Tunable Filter (AA Opto-Electronic, France, AOTFnC-VIS-TN) and triggered with the camera TTL exposure output signal. The laser light is coupled into the microscope by an optical fiber and then reflected using a multi-band dichroic (405 nm/488 nm/561 nm/633 nm quad-band, Semrock, Rochester, NY) and then focused in the back focal plane of the objective. Fluorescence emission light was filtered using a single band-pass filter placed in front of the camera using the following filters: PA-$JF_{549}$: Semrock 593/40 nm bandpass filter; PA-$JF_{646}$: Semrock 676/37 nm bandpass filter. The microscope, cameras, and hardware were controlled through NIS-Elements software (Nikon).

The spaSPT experimental settings for as follows: 1 ms 561nm or 633 nm excitation (100% AOTF) of PA-$JF_{549}$ or PA-$JF_{646}$ was delivered at the beginning of the frame; 405 nm photo-activation pulses were delivered during the camera integration time (~447 μs) to minimize background and their intensity optimized to achieve a mean density of ≤1 molecule per frame per nucleus. 20,000 frames (4 ms) or 30,000 frames (7 or 13 ms) were recorded per cell per experiment. For each cell line or condition, we recorded data for at around 20-30 cells over at least four replicates performed on different days. The camera exposure times were: 4 ms (~223 Hz), 7 ms (~134 Hz), or 13 ms (~74 Hz).

spaSPT data was analyzed (localization and tracking) and converted into trajectories using a custom-written Matlab implementation of the MTT-algorithm ((Sergé et al., 2008); code is available on GitLab: https://gitlab.com/tjian-darzacq-lab/SPT_LocAndTrack) and the following settings: Localization error: $10^{-6.25}$; deflation loops: 0; Blinking (frames): 1; max competitors: 3; max $D$ ($\mu m^2/s$): 20.

#### FRAP

FRAP was performed as previously described (Hansen et al., 2017) on an inverted Zeiss LSM 710 AxioObserver confocal microscope equipped with a motorized stage, a full incubation chamber maintaining 37°C/5% $CO_2$, a heated stage, an X-Cite 120 illumination source as well as several laser lines (only the 561 nm laser was used here). C59 mESCs (Halo-CTCF; Rad21-$SNAP_f$) and C59D2 mESCs (ΔRBR-Halo-CTCF; Rad21-$SNAP_f$) were grown overnight on MatriGel coated glass-bottom (thickness #1.5) 35 mm dishes (MatTak P35G-1.5-14-C), labelled with 500 nM Halo-TMR (Promega Cat. # G8521) for 30 min at 37°C/5.5% $CO_2$ in a TC incubator, washed with PBS and incubated with medium for 5 min in a TC incubator. Cells were then washed again with PBS and the medium changed to medium without phenol red. Images were acquired on a 40x Plan NeoFluar NA1.3 oil-immersion objective at a zoom corresponding to a 100 nm x 100 nm pixel size. 330 frames were recorded at 1 frame every 2 seconds and 20 frames were recorded before the bleach pulse was delivered. A circular bleach spot ($r$ = 10 pixels) was chosen in a region of homogenous fluorescence at a position at least 1 μm from nuclear or nucleolar boundaries. The spot was bleached using maximal laser intensity and pixel dwell time corresponding to a total bleach time of ~1 s. Since the bleach duration was relatively long compared to molecular diffusion, it is not possible to accurately estimate the bound and free fractions from our FRAP curves.

We performed 3 biological replicates recording a total of 18 cells for C59 Halo-CTCF and 18 cells for C59D2 ΔRBR-Halo-CTCF and analyzed the data as previously described (Hansen et al., 2017). As demonstrated previously (Hansen et al., 2017; Sprague et al., 2004), Halo-CTCF and ΔRBR-Halo-CTCF fall in the "reaction dominant" regime, where the recovery depends only on the $k_{\mathrm{OFF}}$ and we therefore fit the FRAP recoveries to the reaction dominant model below:

$$FRAP(t) = 1 - Ae^{-k_a t} - Be^{-k_b t}$$

We interpret the slower rate as specific binding to cognate sites. The fits and this rate are shown in Figure 1-Figure Supplement 1.

#### Data processing and anisotropy calculations

Full details of the data processing and how the calculations were performed including the raw code are available on GitLab: https://gitlab.com/anders.sejr.hansen/anisotropy. Here we will describe each step briefly. We wrote all the code in Matlab and tested it in Matlab 2014b.

First, we converted raw images into SPT trajectories using the MTT-algorithm(Sergé et al., 2008) as described above.

Second, we use "MergeQC_SPT_data.m" to merge data. When performing tracking and localization, each single cell results in a single file. Thus, to keep things manageable, we use "MergeQC_SPT_data.m" to merge data from all these single cells (~20-30 cells) into a single file for a given frame rate, by concatenating the frames (e.g. if two movies with 20,000 frames are merged, frame 1 in the second movie becomes frame 20,001). To minimize tracking errors, we also use "ClosestDist = 2" (in units of μm) to abort trajectories where two particles came closer than 2 μm to each other. This is achieved by calling the function "RemoveAmbigiousTracks.m".

Third, we use "Batch_vbSPT_classify.m" to classify trajectory segments into "bound" and "free" segments. "Batch_vbSPT_classify.m" first removes gaps from all the trajectories and then calls vbSPT(Persson et al., 2013). We implemented a 2-state Hidden Markov Model (HMM) through vbSPT, which uses a Bayesian approach to infer the most likely state for each displacement in a trajectory ("bound" or "free") based on displacement lengths and trajectory history. This is important, because the apparent movement of bound molecules is dominated by localization errors. Thus, we want to exclude bound/immobile molecules from the analysis since we are interested in understanding the nuclear search mechanism, which only applies to diffusing/free molecules and also because bound/immobile molecules will artefactual appear anisotropic due to localization errors around a relatively fixed position. Thus, by using a 2-state HMM we can filter out the bound population and restrict our subsequent analysis to the free population (the number of states is controlled by the variable "maxHidden". "Batch_vbSPT_classify.m" will automatically run on all SPT datasets in the directory "input_path", reformat data to remove gaps and save the reformatted data to "path_reformatted" and finally save the classified data to "path_classified". The final classified SPT data contains 4 variables:

- "CellTracks", a cell array where each element is a trajectory and is a Nx2 matrix with the XY coordinates for each of the N frames.
- "CellTrackViterbiClass", a cell array where each element is a N-1 column vector corresponding to the relevant trajectory in "CellTracks". In other words, "CellTrackViterbiClass" classifies each displacement and thus is one length shorter than the number of localizations in "CellTracks". "1" corresponds to bound and "2" corresponds to free. E.g. if a trajectory had 5 localizations, "CellTrackViterbiClass" will have length 4.
- "vbSPT_metadata", a structure array object containing the most relevant vbSPT metadata such as the inferred diffusion coefficients, subpopulation sizes and transition matrix.
- "LagTime", the time between frames in units of seconds.

"Batch_vbSPT_classify.m" calls two dependent functions: "InferFrameRateFromName.m", which infers the frame rate from the filename and "EditRunInputFile_for_batch.m" which edits the file "vbSPT_RunInputFileBatch.m" to automatically feed the relevant information to vbSPT. In summary, at the end of this step the trajectories have been classified to allow subsequent analysis to focus exclusively on the free/diffusing population.

Fourth, we use "CompileTemporalSubSamplesOfHMM.m" to temporally subsample the existing SPT data and generate trajectories with longer lag times. E.g. by subsampling every $10^{\mathrm{th}}$ frame (frames 1,

11, 21, …) of the 223 Hz data and every 6th frame (frames 1, 7, 13, …) of the 133 Hz data, we can generate new SPT trajectories at 22.2 Hz. Full details on how the 223 Hz, 133 Hz and 74 Hz SPT data was temporally subsampled is given in the structure array, "TempSubSampleStruc" in lines 40-117. We use this approach to generate SPT data at the following frame rates: 44.4 Hz, 34.2 Hz, 26 Hz, 22.2 Hz, 18.8 Hz, 16.5 Hz, 14.8 Hz, 12.2 Hz, 10.6 Hz, 9.2 Hz. The subsampling is performed in the dependent function "TemporallyReSampleCellTracks.m" and we note one potential ambiguity here. To illustrate, suppose we want to take every 3rd frame of a trajectory (i.e. frames 1, 4, 7, …). Since the SPT data has already been HMM-classified, a trajectory of length N will have N-1 displacements classified as either bound or free. We would like to carry over this classification to the temporally subsampled trajectory. While most trajectories are either entirely free ("2") or bound ("1"), some trajectories show transitions. In this example, say the HMM-classification is [1,2,2,2,2,1] for frames 1-7. In this case, the subsampled displacement from frame 1-to-4 will have HMM-classification [1,2,2], but we have to label it as either "1" or "2' in the sub-sampled data. In these cases, we took the most conservative approach. Since our primary goal is to filter out the bound population, we labelled any temporally subsampled displacement as bound as long as any one of the intermediate displacements were classified as bound, even if the majority were free. This is implemented in the function "TemporallyReSampleCellTracks.m". Finally, at the end of this procedure, all the temporally subsampled and HMM-classified SPT datasets are saved to the directory "HMM_first_QC_data".

Fifth, we wrote "Process_SpatioTemporal_AngleAnalysis_v2.m" to perform all the analysis of the HMM-classified trajectory data at multiple spatial and temporal scales. The bulk of the analysis is performed in the dependent function "angleFWHM_Amp_HMM_analyzer_v5.m" and the code also calls "AngleMatrix_analyzer.m" and "ComputeAmpFWHM.m". The analysis is somewhat complicated and for full details, we refer the reader to the underlying function "angleFWHM_Amp_HMM_analyzer_v5.m". But briefly, we describe the analysis, input parameters and output results here. For a given trajectory, as long as it consists of at least 3 localizations and thus 2 displacements, it is possible to calculate an angle between adjacent displacements. If we define the 3 localizations making up the angle as p1, p2 and p3, we can define 2 column vectors and calculate the angle between them as follows (using Matlab syntax):

```
>> v1 = (p2-p1)';
>> v2 = (p3-p2)';
>> angle(1,1) = abs(atan2(det([v1,v2]),dot(v1,v2)));
>> angle(1,2) =  2*pi-angle(1,1);
```

Here, the second element in the angle is due to the factor that whether a 150 degree angle is classified as 150 or 210 degrees is arbitrary. Note though that the above calculation is in units of radians. We then loop over all the trajectories. However, we want to be careful to filter out the bound population and we therefore apply two criteria. First, the displacements must be HMM-classified as "free" and, second, both displacements must be at least of length "MinMinJumpThres". Only if both criteria are satisfied, do we count the angle. These criteria are used for the bulk analysis of the angles (subplots 1-6 in the plotting step). Afterwards, the analysis quantifies 4 different anisotropy metrics:

- AC: anisotropy coefficient which is define as: AC = $\log_2(P(a\in[150°\text{-}210°])/\ P(a\in[330°\text{-}30°]))$, and this metric was introduced previously by Izeddin et al.(Izeddin et al., 2014) Thus, the AC quantifies how much more likely a molecule is to go in the backwards after having going forwards.
- Amp: how the amplitude is calculated is described in the function "ComputeAmpFWHM.m". Briefly, we build a histogram of the probability as a function of the binned angle. We then use fine-scale interpolation to overcome the binning. From this we fit the background density and then calculate the "excess" anisotropy. Since the histogram sums to 1, the amplitude takes values between 0 and 1. This provides a related but somewhat orthogonal metric or anisotropy.
- f(180°±30°/0°±30°) or f(180/0) for short: is identical to the AC, but does not use a logarithm. Essentially, f(180/0) quantifies how many times more likely a particle is to go

backwards relative to continuing forwards. For example, if f(180/0)=1.6, the particle is 1.6-fold more likely to return in the backwards direction than to continue forwards.

- FWHM (full width at half-maximum): quantifies the width of the anisotropy histogram and is defined as the full width of the histogram peak centered around 180° at the half-maximal value (i.e. when the probability is half-way in between the peak of the of the angle histogram and the background probability).

This analysis quantifies the "bulk" anisotropy.

To analyze how the anisotropy changes with space and time, we do binning. For spatial analysis, i.e. how the anisotropy metrics depend on the length of the two displacements making up the angle, we control the range of translocations in the variable "MovingThreshold", which we allow to run from 100 nm to 950 nm in bins of 50 nm. Thus, in this case we additionally consider displacements that are HMM-classified as free and also which are at least 100 nm long (set by the variable "GlotbalMinJumpThres"). Since there are two displacements making up the angle, one can either quantify their mean displacement, their minimal displacement (univariate analysis) or the length of both displacements (bivariate analysis), and we took all three approaches here. For spatial analysis, we averaged over all 13 time-scales (from 223 Hz to 9.2 Hz) and calculated the anisotropy metrics. For example, for f(180/0) as a function of mean displacement, we populated the [100 nm ; 150 nm] bin by taking all angles from all 13 frame rates where the mean of the two displacements making up the angle is between 100 and 150 nm and then calculated f(180/0) using all of these angles. And likewise for the other metrics. For analysis of anisotropy as a function of time, we use all the angles making up a single frame rate satisfying the both of the two criteria listed above: HMM-classified as "free" both displacements at length "MinMinJumpThres" (200 nm).

Errorbars were estimated using re-sampling and is controlled by the variables "JackKnife_fraction" and "JackKnife_iterations". Specifically, we resampled 50% of the angles with replacement 50 times and calculated the anisotropy metrics for each iteration. The error bars reported here show the standard deviation between these 50 iterations.

We noticed that at very long displacements, we would occasionally see very strange trajectories such as a particle shifting back and forth between 2 points separated by a large distance (~800 nm). These are almost certainly a tracking artifact, perhaps from 2 bound molecules blinking out-of-frequency. To remove this and avoid these biasing the analysis especially at long displacements, we removed trajectories where more than half of the angles were highly anisotropic using the variables "MaxNumAngles" and "MaxAsymAnglesFrac".

Finally, all of the anisotropy metrics are saved to the structured array "FinalResults" and saved. With the example data provided on GitLab, the workspace saved is "U2OS_C32_SpatioTemporalAnalysis.mat".

Sixth and finally, we used "PLOT_SpatioTemporalAnalysisResults_v4.m" to plot and visualize the results for each individual data set and similar code to overlay results from multiple different samples. For a more comprehensive description of the analysis and the underlying source code, please see GitLab: https://gitlab.com/anders.sejr.hansen/anisotropy

#### MSD calculations and fitting

Full details and all the raw code used for calculating the time- and ensemble-averaged MSD is available on GitLab https://gitlab.com/anders.sejr.hansen/anisotropy. Here we provide a brief overview. We wrote all the code in Matlab and tested it in Matlab 2014b. The mean-squared displacement (MSD) is a classic analysis approach for SPT. Briefly, it involves increasing the displacements at increasing time-lags and then plotting the mean of the squared distance with time. If the particle exhibits normal Brownian motion, the MSD should be linear with time according to:

$$MSD(\tau) = 4D\tau^{\alpha}$$

Accordingly, α=1 for Brownian motion. α<1 indicates subdiffusion (anomalous diffusion). Here, since our trajectories are generally too short to analyze individually(Michalet and Berglund, 2012), and we therefore compute the time- and ensemble-averaged MSD. The MSD at a timelag τ is given by:

$$MSD_i(\tau) = \big(r_i(t+\tau) - r_i(t)\big)$$

Thus, if we average over all displacements with time-lag τ and over all trajectories, *i*, we obtain the time- and ensemble-averaged MSD for a given timelag τ. In the case of nuclear proteins that bind chromatin and which are tracked with a significant localization error (around 35 nm in our data), the calculations are substantially more complicated. For example, the bound molecules exhibit much longer trajectories than the freely diffusing population, since the free population rapidly moves out-of-focus(Hansen et al., 2018a). Thus, at longer time-lags, unless corrected for, the bound population would dominate the MSD calculations. Moreover, MSDs should not be calculated and averaged over a mixture of two distinct populations. Thus, to filter out the bound molecules, we use the HMM-classification described above. We use the function "MSD_HMM_analyzer.m" to calculate the MSD using only the segments that are classified as free using the HMM. Since we have data at three different frame rates, we calculate the MSD for each frame rate individually (~223 Hz, 134 Hz, 74 Hz) and we also merge the data from all three frame rates.

When it comes to model-fitting, we fit to the HMM-classified MSD. But we must also account for localization errors and we therefore consider the expression:

$$MSD_{\mathrm{HMM}}(\tau) = 4D\tau^{\alpha} + 4\sigma^2$$

where σ is the localization error (standard deviation in one dimension; approximate 35 nm as determined using Spot-On(Hansen et al., 2018a)). In terms of model-fitting the time- and ensemble-averaged MSD, there are a number of considerations. First, how long time-lags to consider for calculating the MSD. Second, what fraction of the data to use for the fitting. To answer the first question, we used subsampling of the data: for each time-lag, we subsampled 50% of the data using 25 iterations and calculated the standard deviation (as shown by the error bars). We then limited the number of timepoints to use (using the vector "Conditions.MSD_timepoints") to 20, 18 and 16 for 223 Hz, 134 Hz and 74 Hz, respectively, such that we did not consider MSD values where the error bars got very large. In terms of least-squares fitting of the data, Saxton has argued that one should limit the fitting to only a fraction of the MSD curve(Saxton, 1997, 2007a). Thus, here we only use the first 50% of the data in the fitting and we perform least-squares fitting as shown in "ProcessPlotFit_MSD.m".

We note that fitting the MSD was not very robust in this case. We see substantially variation in the inferred parameters between the 3 frame rates and it has been argued elsewhere that MSD-analysis is not an optimal method for analyzing SPT data(Lee et al., 2017). We also note that MSD analysis is much less robust for inferring diffusion coefficients(Hansen et al., 2018a). Nevertheless, since it is a standard method for determined the anomalous exponent, α, we include it here. We took the final inferred α as the one fitted when averaged over data from all three frame rates and we also calculated the 95% confidence interval. Full details and raw code to reproduce our analysis is given on Gitlab: https://gitlab.com/anders.sejr.hansen/anisotropy.

#### Brownian motion simulations with simSPT

Even particles obeying ideal Brownian motion can appear anisotropic due to localization uncertainty. For example, a chromatin bound protein subject to 35 nm localization error (defined as Gaussian standard deviation; roughly our experimental localization error) will appear to move around the true position due to the localization uncertainty and this movement will appear highly anisotropic. We therefore filtered out bound molecules as described above (HMM-based removal of the bound population; minimal displacement length for both displacements making up the angle). To validate that our anisotropy pipeline fully filters out spurious apparent anisotropy stemming from localization error, we performed "matched Brownian simulations". Briefly, for each protein and condition, we used Spot-On (Hansen et al., 2018a) to calculated the bound fraction, the free diffusion coefficient and the localization error/uncertainty (~35 nm). We then simulated 500,000 trajectories under realistic HiLo experimental conditions (Figure 1-Figure Supplement 1B) using simSPT ((Hansen et al., 2018a); available on GitLab: https://gitlab.com/tjian-darzacq-lab/simSPT) matching the bound and free diffusion coefficients, the bound fraction and the localization error (35 nm) and simulated data at 223 Hz, 134 Hz, and 74 Hz to match the experiments. We then processed the simulated SPT data identically to how we processed the experimental spaSPT data using the Anisotropy pipeline (available on GitLab:

https://gitlab.com/anders.sejr.hansen/anisotropy). As can clearly be seen in Figure 1-Figure Supplement 1C-D, a combination of 35 nm localization uncertainty and confinement inside the nucleus clearly do contribute mild apparent anisotropy, especially at short time-scales for slow-moving proteins (sim Rad21-Halo; $D_{\text{FREE}}$ = 1.5 μm$^2$/s), but none of the key features observed experimentally such as the "anisotropy bump" around 200 nm are present in the simulations. We conclude that our analysis pipeline successfully filters out apparent anisotropy and that the key experimentally observed features are not artifacts.

#### Calculation of relative ON-rates for wt-CTCF and ΔRBR-CTCF

Here we calculate the effect of the RBR on CTCF's search time within a simplified 2-state model frame-work. Within this simplified 2-state framework the specifically bound fraction for a protein is given by:

$$F_{\text{BOUND}} = \frac{k_{\text{ON}}^*}{k_{\text{ON}}^* + k_{\text{OFF}}}$$

Here, $k_{\text{ON}}^*$ corresponds to the pseudo-first order rate constant and is related to the concentration of available specific binding sites $k_{\text{ON}}^* = k_{\text{ON}}[\text{DNA}]$. The total bound fraction is the sum of non-specific (ns) and specific (s) bound protein: $F_{\text{BOUND,tot}} = F_{\text{BOUND,ns}} + F_{\text{BOUND,s}}$. Previously, we estimated that for CTCF, $F_{\text{BOUND,ns}} = 19.1\%$ (Hansen et al., 2017). Assuming that this is the same for wt-CTCF and ΔRBR-CTCF and noting additionally that $F_{\text{BOUND,tot,wt-CTCF}} = 62.4\%$ (this paper) and $F_{\text{BOUND,tot,}\Delta\text{RBR-CTCF}} = 42.0\%$, we arrive at $F_{\text{BOUND,s,wt-CTCF}} = 43.3\%$ (this number differs by a few percentage points from (Hansen et al., 2017), which is likely due to using a different frame rate and due to experimental variability) and $F_{\text{BOUND,s,}\Delta\text{RBR-CTCF}} = 22.9\%$. Noting that $k_{\text{OFF}}$ does not appear to differ between wt-CTCF and ΔRBR-CTCF (Figure 4, Figure 4-Figure Supplement 1), we can thus calculate the ratio of $k_{\text{ON,wt-CTCF}}^*$ and $k_{\text{ON,}\Delta\text{RBR-CTCF}}^*$.:

$$\frac{k_{\text{ON,wt-CTCF}}^*}{k_{\text{ON,}\Delta\text{RBR-CTCF}}^*} = \frac{\frac{F_{\text{BOUND,s,wt-CTCF}}k_{\text{OFF}}}{1 - F_{\text{BOUND,s,wt-CTCF}}}}{\frac{F_{\text{BOUND,s,}\Delta\text{RBR-CTCF}}k_{\text{OFF}}}{1 - F_{\text{BOUND,s,}\Delta\text{RBR-CTCF}}}} = \frac{\frac{F_{\text{BOUND,s,wt-CTCF}}}{1 - F_{\text{BOUND,s,wt-CTCF}}}}{\frac{F_{\text{BOUND,s,}\Delta\text{RBR-CTCF}}}{1 - F_{\text{BOUND,s,}\Delta\text{RBR-CTCF}}}} = \frac{\frac{0.433}{1-0.433}}{\frac{0.229}{1-0.229}} = 2.57$$

We stress here that this calculation is associated with some uncertainty, since it is difficult to both define and to measure the non-specifically bound fraction. Moreover, the 2-state framework here is somewhat simplified. Nevertheless, given than the specifically bound fraction is substantially higher for wt-CTCF than for ΔRBR-CTCF, the $k_{\text{ON}}^*$ also has to be substantially higher. In other words, the point is not whether the $k_{\text{ON}}^*$ is precisely 2-fold, 2.5-fold or 3-fold higher for wt-CTCF than for ΔRBR-CTCF, but simply that it must be significantly higher.

#### Brownian simulation of chromatin and the protein

To model the dynamics of a protein transiently interacting with chromatin we used the chromatin model developed in (Amitai et al., 2017). The chromatin molecule is represented as a long polymer with $N$ monomer diffusing in a large domain of radius $A$ representing the nucleus. The protein is represented by a diffusing point particle. The chromatin chain is a flexible polymer with spring potential, and Lennard-Jones forces (LJ), describing self-avoidance of each monomer pairs. The polymer has potential energy of the form

$$U(\mathbf{R}_1, ..\mathbf{R}_N) = U_{spring}(\mathbf{R}_1, ..\mathbf{R}_N) + U_{LJ}(\mathbf{R}_1, ..\mathbf{R}_N),$$

where the spring potential is

$$U_{spring}(\mathbf{R}_1, ..\mathbf{R}_N) = \frac{1}{2}k\sum_{i=1}^{N-1}\left(|\mathbf{r}_{i+1,i}| - l_0\right)^2,$$

with $l_0$ is the equilibrium length of a bond, $k = \frac{3}{S_{l_0}^2}$ is the spring coefficient, and $S_{l_0}$ is the standard deviation of the bond length. We chose the empirical relation $S_{l_0} = 0.2l_0$. The Lennard-Jones potential is

$$U_{LJ}^{i,j}(\mathbf{r}_{i,j}) = \begin{cases} 4\left[\left(\frac{\sigma}{\mathbf{r}_{i,j}}\right)^{12} - 2\left(\frac{\sigma}{\mathbf{r}_{i,j}}\right)^{6} + \frac{1}{4}\right] & \text{for} \|\mathbf{r}_{i,j}\| < 2^{1/6}\sigma, \\ 0 & \text{for} \|\mathbf{r}_{i,j}\| \geq 2^{1/6}\sigma \end{cases}$$

where $\sigma$ is the size of the monomer. With the choice $l_0 = 2\sigma$, the springs which materialize bonds, cannot cross each other in stochastic simulations using the potential $U$. We do not account here for bending elasticity. Finally, we used Euler's scheme to generate Brownian simulations. At an impenetrable boundary, each rigid monomer is reflected in the normal direction of the tangent plane.

We recall here the interaction model between the chromatin sites (monomers/zones) and the protein (particle) developed in (Amitai, 2018) (see Figure 2-Figure Supplement 1). The protein diffuses in the nuclear domain until encountering one of the monomer sites by entering the trapping radius $\varepsilon$. The protein is then absorbed at the monomer (zone) with probability $P_{trap}$ or reflected with probability $1 - P_{trap}$. While trapped, the particle is free to diffuse inside the zone which is of radius $\varepsilon$, and is partially reflected from its boundary when it hits it. Every time it hits the boundary it can exit with probability $P_{exit}$. Upon release, the particle is placed at a distance $a$ from absorbing monomer position with the same angular direction from which it escaped and starts diffusing again. Since we take $a > \varepsilon$ and $P_{trap} < 1$, that particle has a finite probability to escape from the absorbing zone rather than rebinding back to it immediately. While inside the zone, the protein can bind at a point inside the zone with Poissonian on-rate $k_{on}$ and then be released with Poissonian off-rate $k_{off}$.

As we observed in the experiment, in some of the protein's trajectory it is bounded at one point (up to our localization error) for the full length of the trajectory. To account for this behavior, we assume that in this state the protein is confined to a small zone, which we denote *cognate binding site* (CBS). Of the total $N$ monomer site, a subset fraction $f_{CBS}$ monomers is chosen randomly as CBSs. We denote other monomer sites as *transiently trapping zones* (TTZ). Hence, the CBS (respectively TTZ) has trapping radius $\varepsilon_{CBS}$ (respectively $\varepsilon_{TTZ}$), release radius $a_{CBS}$ (respectively $a_{TTZ}$), capture probability $P_{abs,CBS}$ (respectively $P_{abs,TTZ}$). The protein has a characteristic binding time $\tau_{CBS}$ within the CBSs. The third kind of monomers are the Power-law-distributed-Trapping Zones (PTZs). Each has a different size which is drawn out of a power-law distribution $P_{PTZ}(\varepsilon)$ with a size cutoff at 800nm. The probability of binding to a PTZ upon encountering one is inverse to its size ($P_{trap,PTZ} = \frac{A_1}{\varepsilon^{\delta}}$). Inside the TTZs and the PTZs, the protein diffused and is partially reflected from its boundary when it hits it. The exit probability $P_{exit}$ is the same for TTZs and PTZs. The release radius of each PTZs is different and is larger by 10nm from its trapping radius $\varepsilon$.

At the equilibration stage of the system, the polymer is placed inside the nuclear domain and equilibrates for a time T longer than its longest relaxation mode. The end monomer remains fixed at the origin. After this initial phase, the polymer configuration is fixed. The protein is then placed at a random position in the nuclear domain. The protein position evolves in time according to the Langevin equation

$$dx = \sqrt{2\mathrm{D}_i}\, d\omega, \qquad i = F, Z, CBS$$

where $D_Z$ is the diffusion coefficient of the protein within a zone, $D_F$ is the diffusion coefficient a free protein (outside of the zones), $D_{CBS}$ is the diffusion coefficient within the small CBS, and $d\omega$ is a white Gaussian noise. The protein diffuses until encountering a trapping zone or a CBS as detailed above. For each unique polymer configuration for simulated the protein motion for 10000 absorption events. In each condition, we randomized many different polymer configurations.

Each simulation produced a long Brownian trajectory of the protein in the nuclear domain. White Gaussian noise with standard deviation of 30nm and mean zero is added to each trajectory point. The trajectories from the simulations were aggregated and then split to short trajectories with the same length statistics as that of the experimental trajectories. The trajectories were then subjected to the same classification procedure by an HMM as were the experimental trajectories. We then computed different statistics of these trajectories.

#### APPENDIX 1 - Estimating the contribution of tracking errors (misconnections)

Single-particle tracking (SPT) is uniquely suited to analyzing the dynamics of proteins, how they explore the nucleus and whether they exhibit anomalous diffusion since it allows direct observation of molecular motion at high spatiotemporal resolution. However, experimental SPT is also subject to numerous biases, which complicates the interpretation of SPT experiments. First, while a frame is recorded, fast-diffusing molecules can move several pixels, which causes a "motion-blur" artifact. These molecules will not be recognized by most localization algorithm since they no longer resemble a PSF and thus fast-diffusing molecules will be undercounted. Second, most SPT experiments in live mammalian cells have used quite high particle densities (e.g. ~20-100 in-focus molecules per frame). This allows a user to rapidly and conveniently record large amounts of data, but comes at the cost of tracking errors. When the displacements of multiple particles overlap between frames, there is no way to unambiguously identify which molecule moved where. In general, the fraction of displacements that are misconnected increases with particle density. In the high-density limit, when the trajectories contain many tracking errors, it is no longer possible to make inferences about how single molecules explore the nucleus.

To minimize these biases we introduced stroboscopic photo-activation single-particle tracking (spaSPT)(Hansen et al., 2017, 2018a), which builds on and integrates previous approaches (Elf et al., 2007; Manley et al., 2008). First, spaSPT minimizes motion-blur bias by using stroboscopic excitation (Elf et al., 2007). Here we use 1 ms stroboscopic excitation pulse, which is sufficient to achieve very good signal (signal-to-background ratio > 5), and which we have previously shown theoretically (Hansen et al., 2017) and experimentally (Hansen et al., 2017, 2018a) causes negligible motion-blur bias even for fast-diffusing proteins such as Halo-3xNLS ($D_{\mathrm{FREE}}$ ~ 10-12 μm$^2$/s). Second, spaSPT largely avoids tracking errors, by keeping the particle density very low. We take advantage of the very bright photo-activatable Janelia Fluor dyes (Grimm et al., 2016), which allows us to tune the photo-activation probability such that essentially any desired particle density can be achieved. Here we aimed for an average density of ~ 0.5-1 localizations per nucleus per frame, which provides a reasonable compromise between minimizing tracking errors and also recording large SPT datasets necessary for statistical analysis.

Even though tracking errors through misconnections between 2 particles are quite rare in spaSPT at low densities, they always will happen with some probability and we wanted to quantitatively assess their frequency. To do so, we labeled Halo-tagged proteins in live U2OS cells with the two spectrally distinct dyes, PA-JF$_{646}$ and PA-JF$_{549}$. The HaloTag protein binds only one dye covalently and irreversibly and under our imaging conditions there was no spectral overlap (i.e. any fluorescence bleedthrough between the color channels was far below the detection limit). Thus, we recorded simultaneous 2-color spaSPT data using U2OS C32 Halo-CTCF and U2OS Halo-3xNLS at 133 Hz and 74 Hz. We considered both Halo-CTCF and Halo-3xNLS since they represent opposite extremes: U2OS Halo-CTCF diffuses very slowly ($D_{\mathrm{FREE}}$ ~ 2.0-2.5 μm$^2$/s) and exhibits a high bound fraction ($F_{\mathrm{BOUND,total}}$ ~ 60%), whereas Halo-3xNLS diffuses extremely rapidly ($D_{\mathrm{FREE}}$ ~ 10-12 μm$^2$/s) and exhibits a minimal bound fraction ($F_{\mathrm{BOUND}}$ ~ 10%). Thus, we reasoned that considering the two extremes would allow us to place upper and lower bounds and the frequency of tracking errors.

We then analyzed the data as follows: We analyzed the JF$_{549}$ and JF$_{646}$ datasets individually and we then combined the localizations for both datasets and feed them into the tracking algorithm (Sergé et al., 2008) such that the tracking algorithm was blind to a particle's "color" (and keeping all algorithm parameters the same). This allowed us to identify any cases where a single-trajectory contains localizations in more than one color, which must have involved a tracking misconnection. Moreover, to generate a "worst-case scenario" dataset, we used a particle density of ~0.5-1.5 for JF$_{549}$ and JF$_{646}$ individually (so ~1-3 in-focus particles per frame in total), such that the merged localizations would exhibit a 2-3 fold higher density than normal. Thus, this analysis would generate an upper bound on the number of tracking errors.

However, not all tracking errors will give a change in particle colors. Even in the 2-color datasets, JF$_{549}$-to-JF$_{549}$ and JF$_{646}$-to-JF$_{646}$ are also possible. More generally, if we refer to the colors as "1" and "2", the fractions of all localizations in each color will be:

$$f_1 = \frac{n_1}{n_1+n_2} \text{ and } f_2 = \frac{n_2}{n_1+n_2}$$

where $n$ is the number of localizations per color per movie. If tracking is totally arbitrary (this is a slight approximation, since the probability of either color appearing in frame $k$+1 is not independent of frame $k$ due to photo-activation being less likely than photo-bleaching), the probability of tracking being observed as correct is:

$$P_{\text{correct}} \approx P(c_{\text{Frame k+1}} = 1 | c_{\text{Frame k}} = 1) + P(c_{\text{Frame k+1}} = 2 | c_{\text{Frame k}} = 2)$$

and similarly the probability of actually observing a tracking error is:

$$P_{\text{wrong}} \approx P(c_{\text{Frame k+1}} = 1 | c_{\text{Frame k}} = 2) + P(c_{\text{Frame k+1}} = 2 | c_{\text{Frame k}} = 1)$$

Again, if we assume tracking to be totally arbitrary and that there is no memory between frames (which is an approximation), these reduce to:

$$P_{\text{correct}} \approx P(c_{\text{Frame k+1}} = 1 | c_{\text{Frame k}} = 1) + P(c_{\text{Frame k+1}} = 2 | c_{\text{Frame k}} = 2)$$
$$= f_1 \cdot f_1 + f_2 \cdot f_2 = {f_1}^2 + (1 - f_1)^2 = 1 + 2({f_1}^2 - f_1)$$

And

$$P_{\text{wrong}} \approx P(c_{\text{Frame k+1}} = 1 | c_{\text{Frame k}} = 2) + P(c_{\text{Frame k+1}} = 2 | c_{\text{Frame k}} = 1)$$
$$= f_1 \cdot f_2 + f_2 \cdot f_1 = 2f_1(1 - f_1) = 2(f_1 - {f_1}^2)$$

Thus, as expected in the limit where tracking is totally arbitrary and where $f_1 = f_2$, these reduce to $P_{\text{correct}} \approx 0.5$ and $P_{\text{wrong}} \approx 0.5$. In practice, due to slight differences in cell permeability and labeling kinetics between $JF_{549}$ and $JF_{646}$ (Yoon et al., 2016), as well as experimental variations such as small pipetting errors and slight differences in photo-stability etc., it is difficult to get exactly 1:1 labeling and precisely the same number of localizations for both dyes. While the exact expressions above can be used for individual movies based on the number of localizations for each dye, in order to get good statistics it is necessary to averages over many movies with slightly different fractions. Moreover, in almost all of the movies the fraction of localizations in one color was in the range 40-60%. $P_{\text{wrong}}$ is symmetric and inverse U-shaped, and even $P_{\text{wrong}}(f_1 = 0.4) = P_{\text{wrong}}(f_1 = 0.6) = 0.48$, which is very close to 0.5, as expected if we obtained perfectly 1:1 labeling. Thus, averaging over all movies despite these having slightly different relative numbers of localizations is a very small approximation.

Having verified that we can reasonably average all of the movies, we next analyzed how the number of observed tracking errors depends on density and displacement length. First, we note that the probability of a tracking error resulting in a color change is approximately 50% assuming $f_1 \approx f_2$. That is, if we are tracking a particle in color 1, a misconnection to another particle in color 1 will not appear as a tracking error since it does not result in a color change. Only if the misconnection occurred to a particle of the opposite color, will it appear as a color change. Thus, when we count tracking errors as color changes, the actual number of tracking errors is ~2-fold higher than the observed number assuming $f_1 \approx f_2$. Thus, we multiplied the observed number of tracking errors by 2 to get the total number of tracking errors, both observed and unobserved. Somewhat encouragingly, after multiplying by 2 the fraction of incorrect connections, it was still quite low despite imaging at a 2-3 fold higher particle density than normal. Specifically, we got the following number of incorrect connections (after multiplying by 2): U2OS C32 Halo-hCTCF; 74 Hz gave 3.36%; U2OS C32 Halo-hCTCF; 133 Hz gave 1.85%; U2OS Halo-3xNLS; 74 Hz gave 6.65%; U2OS Halo-3xNLS; 133 Hz gave 4.12%.

As expected, the number of tracking errors increase when the frame rate decreases (74 Hz vs. 133 Hz) since the mean displacement increases with the lag time and also as expected, the number of tracking errors is higher for Halo-3xNLS, which diffuses very rapidly (mean displacement at 74 Hz: 637 nm) and thus is more likely to cross paths with another molecule than it is for Halo-hCTCF (mean displacement at 74 Hz: 178 nm), which diffuses much more slowly. Thus, even at 2-3 fold higher particle density, the number of tracking errors for Halo-hCTCF at 133 Hz is still <2%.

We further analyzed the 2-color spaSPT data to understand under which conditions tracking errors start appearing (Figure 1-Figure Supplement 2A). This analysis reveals several important points: First, the mean displacement for misconnections is much longer than for correct tracking connections (e.g. 134 nm vs. 766 nm for Halo-hCTCF at 133 Hz). Second, although the mean number of localizations per frame is generally higher when there is a misconnection compared to when the tracking was correct, it is only slightly higher (e.g. 2.05 localizations/frame for correct connections vs. 2.35 localizations/frame for misconnections for Halo-hCTCF at 133 Hz; please note that this is not the average number of localizations per frame, which is significantly lower; but since we only count displacements, e.g. frames without localizations are not counted). Third, the probability of a displacement being a misconnection (tracking error) increases *exponentially* with the displacement length (until saturation). This is perhaps surprising at first glance and to understand why this would be and to facilitate filtering out misconnections, we will distinguish two types of tracking errors: Misconnection due to Overlapping Trajectories (MOT) and Misconnection due to new Appearance of Particle (MAP) (Figure 1-Figure Supplement 2B).

Misconnection due to Overlapping Trajectories (MOT; Figure 1-Figure Supplement 2B): MOT errors will tend to occur at high densities. The higher the number of particles, the more ambiguous the tracking becomes. While many tracking algorithms (Lee et al., 2017), including the MTT-algorithm (Sergé et al., 2008) used here, are more sophisticated than simply connecting nearest neighbors (for example by taking trajectory history into account), as trajectories begin to overlap, there is simply no way of unambiguously connecting molecules between frames. Accordingly, we expect the number of MOT errors to increase with: 1) increasing density of particles; 2) longer lag times (i.e. time between frames; slower frame rates); 3) larger molecular diffusion coefficient (e.g. since CTCF diffuses more slowly than Halo-3xNLS ($D_{\mathrm{FREE}}$ ~ 2.5 μm$^2$/s vs. 10-12 μm$^2$/s) trajectories are less likely to overlap for a given density and frame rate since the average displacement is much lower; accordingly, the fraction of misconnections is ~2-fold smaller for CTCF compared to Halo-3xNLS).

Misconnection due to new Appearance of Particle (MAP; Figure 1-Figure Supplement 2B): MAP errors tend to occur when a new particle appears close to an existing particle. This can either be due to a particle outside of the focal plane moving inside the focal plane (this may occur regardless of whether photo-activation is used or not) or due to spontaneous photo-activation of a particle close an existing particle. If the existing particle either bleached or moved further away, a misconnection to the newly appeared particle is then likely. MAP errors due to particles out-of-focus moving into focus are density dependent, but depend on the density of out-of-focus particles and thus will not necessarily be captured in a simple statistic like localizations/frame, which only count in-focus and detected localizations. Conversely, MAP errors due to photo-activation might be expected to be inversely dependent on density: the probability of such a MAP error occurring increases at low densities where any newly photo-activated molecule will be connected to a molecule that photobleached in the previous frame assuming that the two were sufficiently close. It is also important to note that MAP errors due to photo-activation should be expected to scale with the displacement squared. If we assume that photo-activation is equally likely to occur anywhere in the nucleus, the probability of such a MAP error to occur will depend on the area of the maximally allowed displacement ($A(r) = \pi r^2$). Thus, for this reason, we might initially expect the misconnection probability to increase with $r^2$ instead of exponentially as observed above. However, theory actually predicts the misconnection probability to increase exponentially with distance. To understand why, consider a case of a protein (very similar to CTCF) where roughly half of the molecules are bound to chromatin and with $D_{\mathrm{FREE}}$ ~ 2.5 μm$^2$/s. The distribution of displacements (leaving out the normalization constant) is then given by:

$$P(r) = F_{\mathrm{BOUND}} \frac{r}{2(D_{\mathrm{BOUND}}\Delta\tau + \sigma^2)} e^{-\frac{r^2}{4(D_{\mathrm{BOUND}}\Delta\tau + \sigma^2)}} + (1 - F_{\mathrm{BOUND}}) \frac{r}{2(D_{\mathrm{FREE}}\Delta\tau + \sigma^2)} e^{-\frac{r^2}{4(D_{\mathrm{FREE}}\Delta\tau + \sigma^2)}}$$

Here, $F_{\mathrm{BOUND}}$ is the fraction of molecules that are bound to chromatin, $D_{\mathrm{BOUND}}$ is diffusion coefficient of chromatin bound molecules, $D_{\mathrm{FREE}}$ is diffusion coefficient of freely diffusing molecules, $r$ is the displacement length, $\Delta\tau$ is lag time between frames and $\sigma$ is localization error. If we assume that MAP errors contribute to a 5% misconnection probability at 500 nm as seen in the raw 2-color Halo-hCTCF data at 134 Hz such that $A(r = 500\ \mathrm{nm}) = 0.05P(r = 500\ \mathrm{nm})$, then we get the results shown in Figure 1-Figure Supplement 2C. On the top plot, we see that at long displacements, MAP errors (red) become more likely than true displacements (green). If we now plot the fraction of displacements that are incorrect as a fraction of all displacements (right plot), we see precisely the same trend as in the experimental data: We see a steep exponential increase at lower displacements (up until ~100-150 nm), followed by a kink and then a slower exponential increase from around ~150 nm to ~700 nm, where saturation begins to happen. The misconnection probability increases exponentially because the correct connection probability decreases exponentially and thus the exponential trend is due to $P(r)$ and not $A(r)$. Of course, where the transitions occur depend on the details of the parameters and this model involves several approximations, but it is notable that such a simple theoretical model can faithfully capture observed experimental trends. Most importantly, it highlights that tracking errors increase exponentially with displacement length.

The above analysis revealed MOT errors to be highly density dependent and MAP errors somewhat density dependent. Consistently, misconnections occur at, on average, slightly higher particle densities (Figure 1-Figure Supplement 2A). We note that although the above 2-color experiments were performed at a 2-3-fold higher particle density than the 1-color experiments and thus represent a worst-case scenario, occasionally frames with high densities are unavoidable. Since photo-activation and photo-bleaching are both Poisson processes, we would expect the distribution of localizations per frame to roughly follow a Poisson distribution. For example, if the mean density is 1 localization/frame, 26.4% of frames will nevertheless have 2 or more localizations. Likewise, even at a very low mean density of 0.5 localizations/frame, 9.0% of frames will have 2 or more localizations. Thus, a simple approach to minimizing MOT errors is to abort and remove all parts in frames n+1 and beyond, if two trajectories get closer than a certain threshold distance, $r_{\text{threshold}}$. To determine how to choose $r_{\text{threshold}}$ we plotted the probability of misconnections as a function of $r_{\text{threshold}}$ (Figure 1-Figure Supplement 2D). A couple of points stand out: First, as expected, restricting the closest allowable distance between particles in the same frame reduces the number of tracking errors, i.e. the probability of a misconnection. Second, the strategy is effective both for Halo-CTCF (slow diffusion, overwhelmingly bound) and Halo-3xNLS (very fast diffusion, overwhelmingly free). E.g. setting $r_{\text{threshold}} = 2\ \mu\text{m}$, reduces the misconnection probability from 6.77% to 4.55% for Halo-3xNLS at 74 Hz but also from 1.85% to 0.94% for Halo-CTCF at 133 Hz. Third, in all four cases diminishing returns are observed and eventually the curve plateaus. This happens 1.5 μm for Halo-CTCF at 133 Hz and around 2 μm in the other three cases. Most likely, the plateau occurs because this approach can filter out MOT, but not most MAP, errors. Thus, it is likely that the plateau is largely composed of MAP errors.

Importantly, filtering out data in this way also filters out some correctly tracked particles and there is always the risk of introducing unintended biases. Therefore, using $r_{\text{threshold}} = 2\ \mu\text{m}$ seemed like the optimal choice and we therefore applied this to all the datasets ("ClosestDist" variable in MergeQC_SPT_data.m).

Analyzing anisotropy at multiple spatiotemporal scales has proven very informative in distinguishing different mechanistic models of anomalous diffusion (Izeddin et al., 2014). Thus, we would like to determine how far out we can extend the analysis in space (i.e. displacement length) for a given time (i.e. frame rate), while maintaining a relatively low probability of misconnections. We therefore applied the $r_{\text{threshold}} = 2\ \mu\text{m}$ to the raw 2-color dataset and then sampled the trajectories at different time scales. For example, considering localizations 1, 3, 5, 7… of trajectories recorded at 74 Hz, yields a dataset at 37 Hz. Since photobleaching is quite high under our spaSPT conditions, re-sampling the data in this way leads to smaller datasets. At 12.2 Hz, we only had 7,805 Halo-3xNLS trajectories and 32,563 Halo-CTCF trajectories. At this point, analyzing the as above by binning based on displacement length etc. causes the analysis to start to get noisy due to the limited amount of data, so we did not further extend the analysis to longer lag times. First, we plotted the probability of misconnections (after applying $r_{\text{threshold}} = 2\ \mu\text{m}$) as a function of the frame rate or lag time (Figure 1-Figure Supplement 2E). As observed before, Halo-CTCF had fewer tracking errors than Halo-3xNLS and the 133 Hz data had fewer tracking errors than the 74 Hz data (this is why the trend is slightly irregular, since the second point is composed exclusively of 74 Hz data, whereas the third point is composed exclusively of 133 Hz data). Nevertheless, the probability of misconnections did not increase with increasing lag time – if anything, it slightly decreased. Thus, this suggests that we can extend our analysis to substantially longer lag times than under which the data was originally recorded – for example, for Halo-CTCF at ~12.2Hz the probability of a tracking error is just 0.58%, that is, just 1 out of every 172 tracking connections was an error.

To further understand how tracking errors occurred in the temporally subsampled datasets, we repeated the analysis above for both Halo-CTCF and Halo-3xNLS at two representative frame rates: 35

Hz and 12.2 Hz (Figure 1-Figure Supplement 2F). As can be seen, most of the conclusions derived from the analysis of the 74 Hz and 133 Hz datasets, also apply to the temporally subsampled datasets (shown: 35 Hz and 12.2 Hz). In particular, the probability of a displacement being a misconnection reaches our somewhat arbitrary 5%-threshold once we reach displacements of ~1000 nm. Thus, this analysis suggests that we can reliably only analyze displacements up to about ~1000 nm, even at longer timescales if we want to make sure that we are not affected by tracking errors. Also, note that since the trajectories are quite short, when we temporally subsample very long lag times, the amount of data available decreases a lot, which is why the plots in Figure 1-Figure Supplement 2F are more noisy.

In summary, the above analysis and Figure 1-Figure Supplement 2 show that tracking errors contribute minimally to our results as long as we do not consider displacements that are much longer than 800 nm.

### Figure Supplements and Legends and Supplementary Table 1

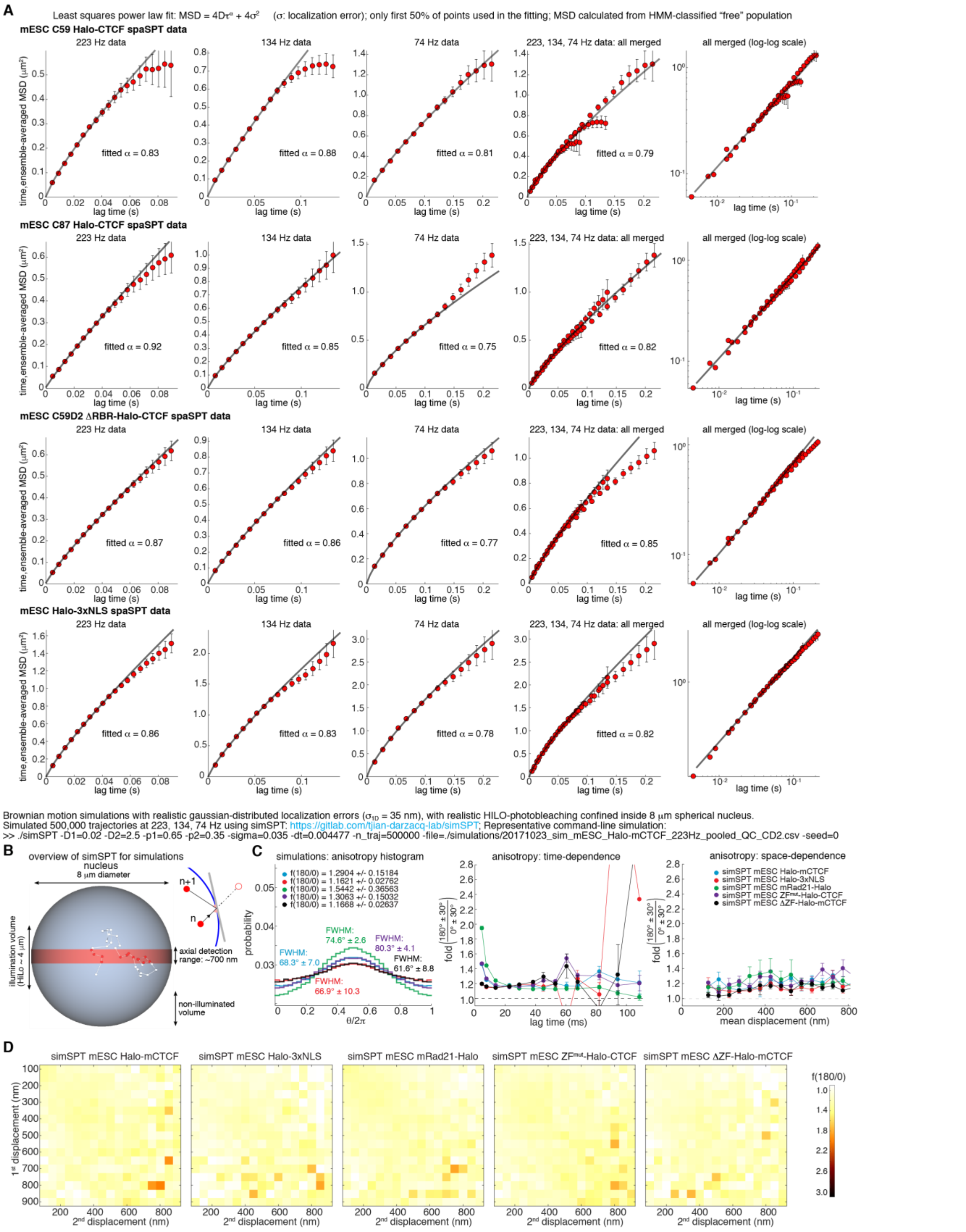


**Figure 1-Figure Supplement 1. MSD-analysis and control Brownian motion simulations.**
(**A**) Mean-square displacement (MSD) analysis for mESC C59 Halo-CTCF (top row), mESC C87 Halo-CTCF (2nd row), mESC C59D2 ΔRBR-Halo-CTCF (3rd row) and mESC Halo-3xNLS (4th/bottom row). The data for each

cell line was analyzed using code available at https://gitlab.com/anders.sejr.hansen/anisotropy. Briefly, using an HMM we filter out the bound population, such that the MSD is only calculated for the diffusing subpopulation. spaSPT (Hansen et al., 2017, 2018b) was performed at three frame rates (4.477 ms per frame or ~223 Hz, 7.477 ms per frame or ~134, and 13.477 ms per frame or ~74 Hz) and the MSD-fit is shown to each of the three datasets in the first three columns. Briefly, a power-law accounting for localization error was fit and the anomalous exponent, $\alpha$, inferred. As suggested (Saxton, 1997, 2007b), we only used the first 50% of the data for the fitting. Error bars show standard deviation (from subsampling 50% of the data using 25 iterations) and the number of timepoints was limited to 20, 18, and 16 for 223 Hz, 134 Hz, and 74 Hz, respectively. The 4$^{th}$ column shows the MSD-fit with all the data merged. We show all four versions in part to emphasize that the inferred anomalous exponent, $\alpha$, is quite sensitive to the data processing and therefore should be interpreted cautiously. Nevertheless, we believe the merged fit (column 4) most robustly represents the actual value. Finally, the 5$^{th}$ column shows the same data at a log-scale.

(**B**) simSPT simulations. Brownian motion with a mixture of bound and free subpopulations subject to significant localization error (std=35 nm) confined inside the nucleus can result in "apparent" anisotropy, since a bound molecule will appear to take steps back and forth around its true localization because of localization uncertainty. The purpose of these simulations was to assess whether or not our analysis pipeline fully filters this out. We performed these simulations using simSPT (Hansen et al., 2018b), which simulations experimentally realistic HILO SPT data. For each condition (e.g. C59 Halo-CTCF at 223 Hz), we "matched" the parameters ($F_{BOUND}$, $D_{FREE}$, $\sigma$) to the data (parameters obtained from analysis with Spot-On (Hansen et al., 2018b)) and then simulated 500,000 trajectories and analyzed them the same was as the experimental data.

(**C**) Analysis of simSPT simulations. Representative plots of fold-anisotropy, $f_{180/0}$, at the bulk level (left), as a function of the lag time (middle) and as a function of the mean displacement length (right) for the indicated "matched" experiments: mESC Halo-CTCF ($F_{BOUND}$=0.65; $D_{FREE}$=2.5 $\mu m^2/s$), mESC Halo-3xNLS ($F_{BOUND}$=0.10; $D_{FREE}$=6.0 $\mu m^2/s$), mESC Rad21-Halo G1 ($F_{BOUND}$=0.50; $D_{FREE}$=1.5 $\mu m^2/s$), mESC ZF$^{mut}$-Halo-CTCF ($F_{BOUND}$=0.30; $D_{FREE}$=2.5 $\mu m^2/s$), mESC ΔZF-Halo-CTCF ($F_{BOUND}$=0.05; $D_{FREE}$=5.5 $\mu m^2/s$). As can be seen, simulated Rad21 is mildly anisotropic at ~223 Hz, but otherwise, the analysis pipeline successfully filters out "apparent" anisotropy.

(**D**) Anisotropy heatmaps. Plot of $f_{180/0}$ as a function of the length of the first and second displacement. Most notably, the anisotropy "bump" around 200 nm is fully absent in the simulations.

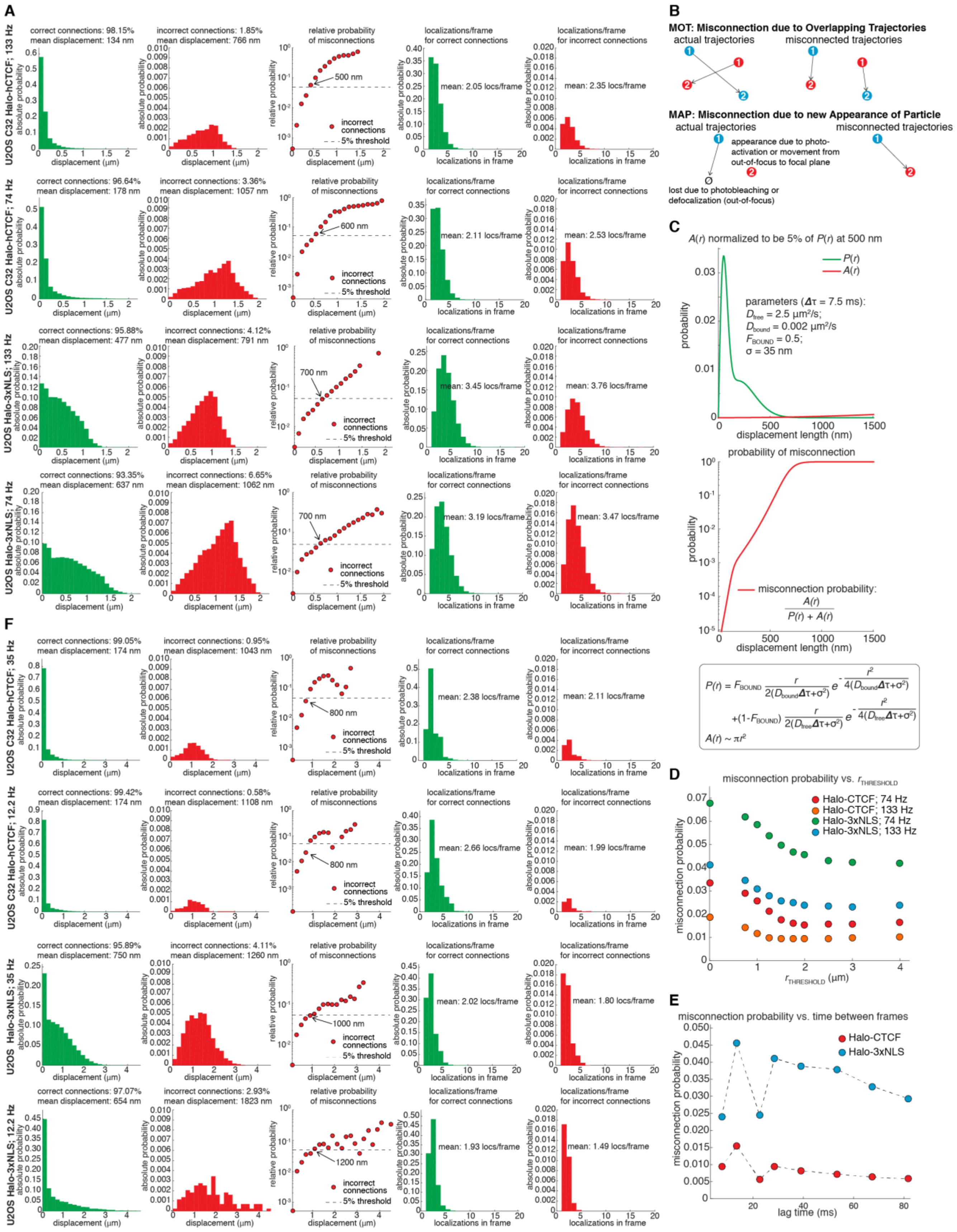


$$P(r) = F_{BOUND} \frac{r}{2(D_{bound}\Delta\tau+\sigma^2)} e^{-\frac{r^2}{4(D_{bound}\Delta\tau+\sigma^2)}} + (1-F_{BOUND}) \frac{r}{2(D_{free}\Delta\tau+\sigma^2)} e^{-\frac{r^2}{4(D_{free}\Delta\tau+\sigma^2)}}$$

$$A(r) \sim \pi r^2$$

**Figure 1-Figure Supplement 2. 2-color spaSPT experiments to estimate the misconnection probability during tracking.**

(**A**) Misconnection probability for U2OS Halo-CTCF and U2OS Halo-3xNLS. Cells were labeled ~1:1 with the distinguishable PA-JF549 and PA-JF646 dyes (Grimm et al., 2016). This way, we can directly observe tracking misconnections (e.g. green-to-red or red-to-green). We correct for the ~2-fold undercounting. We imaged at a

~2-3-fold higher density than normal (~2-3 particles per frame per nucleus) to estimate the worst-case level of tracking errors. Histograms of displacement lengths are shown for correct connections (1st column) and misconnections (2nd column). The 3rd column shows that the misconnection probability increases *exponentially* with displacement length. Histograms of number of localizations per frame for correct and incorrect connections are shown in column 4 and 5, respectively.

(**B**) Schematics. Schematics illustrating two types of misconnections: Misconnection due to overlapping trajectories (MOT) and Misconnection due to new Appearance of Particle (MAP).

(**C**) Why tracking errors increase exponentially. "back-of-the-envelope" illustration of why tracking errors increase exponentially with displacement length. Allow the misconnection probability presumably scales with the area of a circle (radius square), because correct connections decrease exponentially with displacement length, misconnection probability will increase exponentially with displacement length.

(**D**) Close encounter filter. To minimize misconnections, we applied a close encounter filter, $r_{THRESHOLD}$, such that whenever two particles come closer than $r_{THRESHOLD}$ they will be cut short. Choosing $r_{THRESHOLD}$ =2 μm optimally reduced the misconnection probability.

(**E**) How misconnections scale with lag time.

(**F**) Misconnection probability for U2OS Halo-CTCF and U2OS Halo-3xNLS. Same as (**A**), but for temporally subsampled data (35 Hz, 12.2 Hz).

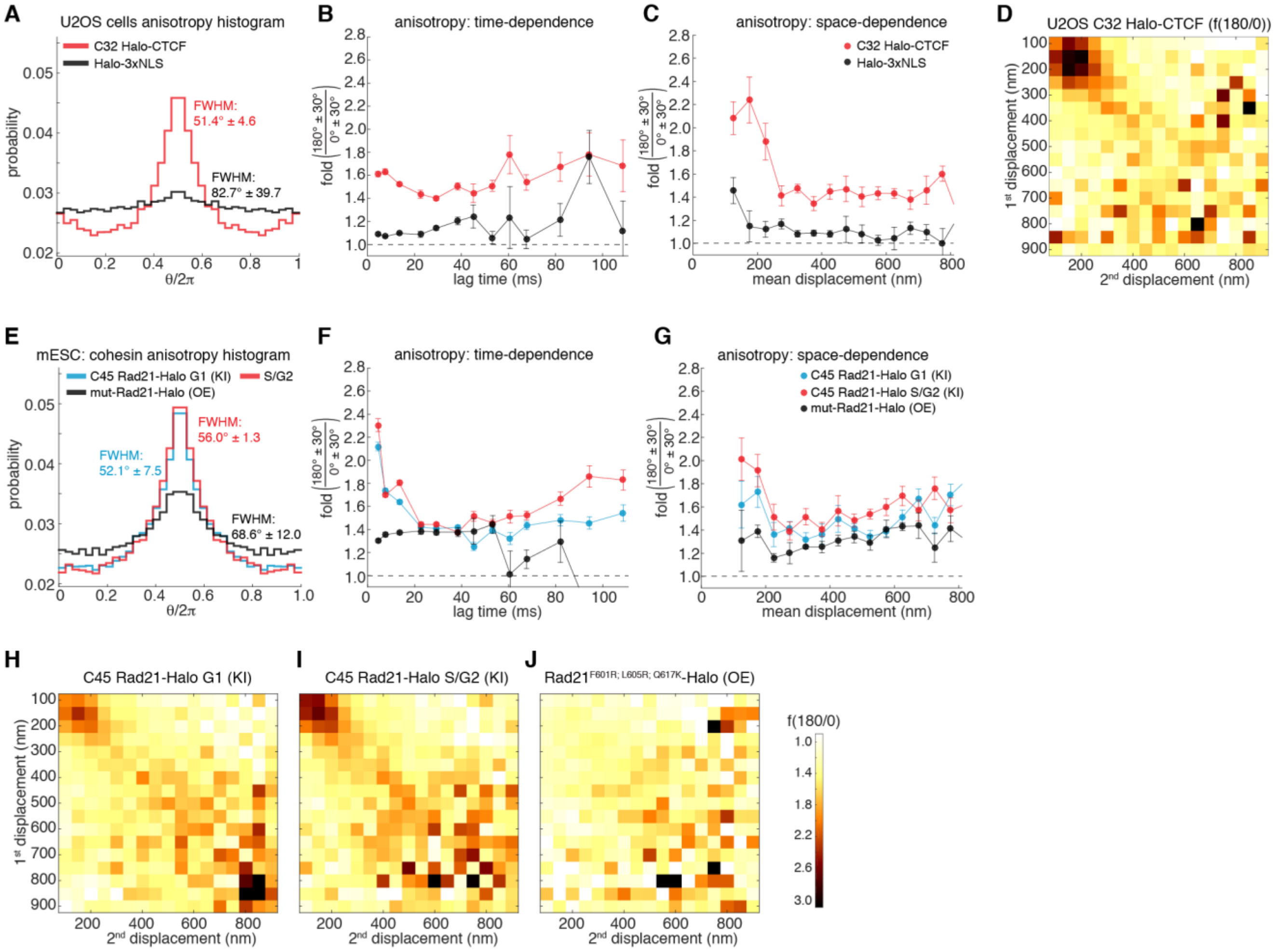


**Figure 1-Figure Supplement 3. Anisotropy for CTCF in human U2OS cells and cohesin in mESCs.**
Anisotropy plots for C32 Halo-CTCF and Halo-3xNLS in human U2OS cells showing fold-anisotropy, $f_{180/0}$, at the bulk level (**A**), as a function of the lag time (**B**) and as a function of the mean displacement length (**C**). (**D**) Anisotropy heatmap. Plot of $f_{180/0}$ as a function of the length of the first and second displacement for U2OS C32 Halo-CTCF.

Anisotropy plots for C45 Rad21-Halo (knock-in) in either G1-phase or S/G2 phase of the cell cycle, and a mut-Rad21-Halo, which cannot form cohesin complexes (Hansen et al., 2017), (transiently expressed; ΔZF10,11-ΔRBR-Halo-CTCF) in mESCs showing fold-anisotropy, $f_{180/0}$, at the bulk level (**E**), as a function of the lag time (**F**) and as a function of the mean displacement length (**G**). (**H-J**) Anisotropy heatmaps. Plot of $f_{180/0}$ as a function of the length of the first and second displacement for the indicated cell lines.

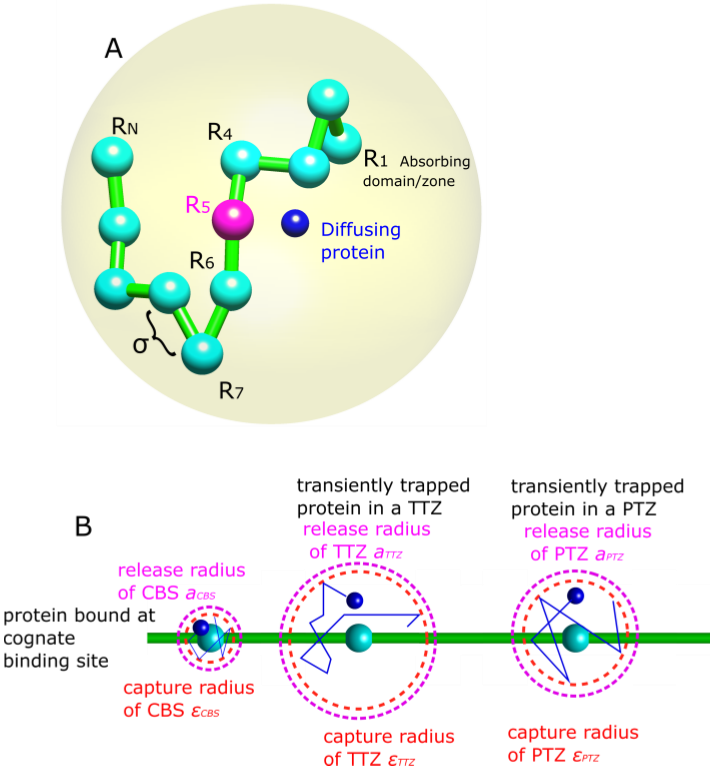


**Figure 2-Figure Supplement 1. Overview of the model.** (**A**) Chromatin is described as a long chain of N monomers (beads, cyan) connected by harmonic springs. The monomers additionally interact via LJ interactions with a LJ size σ. Each monomer corresponds to one of three zones/domains (TTZs,CBSs,PTZs). The monomer assignment is random. The polymer is equilibrated within a domain (nucleus) which has a radius of 5μm. The CTCF protein is a Brownian particle (dark blue). The protein was released from monomer (zone) 5 after being trapped there (magenta). (**B**) (**Left)** The protein is absorbed/trapped with probability $P_{trap,CBS}$ if it arrive to a distance $\varepsilon_{CBS}$ (red dashed-line circle) from the center of a cognate binding site (CBS). It is then trapped there for a characteristic time $\tau_{\text{CBS}}$. When released, the protein is positioned with equal probability on a sphere center around the CBS with a radius $a_{CBS}$ (magenta dashed-line circle). (**Middle**) The protein is absorbed/trapped with probability $P_{trap,TTZ}$ if it arrive to a distance $\varepsilon_{TTZ}$ (red dashed-line circle) from the center of a Transiently Trapping Zone (TTZ). It diffuses within the TTZ which has radius $\varepsilon_{TTZ}$. When released, the protein is positioned on a sphere center around the CBS with a radius $a_{TTZ}$, at the same angular direction from which it escaped (magenta dashed-line circle). (**Right**) The protein is absorbed/trapped with probability $P_{trap,PTZ}$ if it arrive to a distance $\varepsilon_{PTZ}$ (red dashed-line circle) from the center of a Transiently Trapping Zone (PTZ). The distance $\varepsilon_{PTZ}$ is randomized from a distribution (see Material and Methods). It diffuses within the TTZ which has radius $\varepsilon_{PTZ}$. When released, the protein is positioned on a sphere center around the CBS with a radius $a_{PTZ}$, at the same angular direction from which it escaped (magenta dashed-line circle).

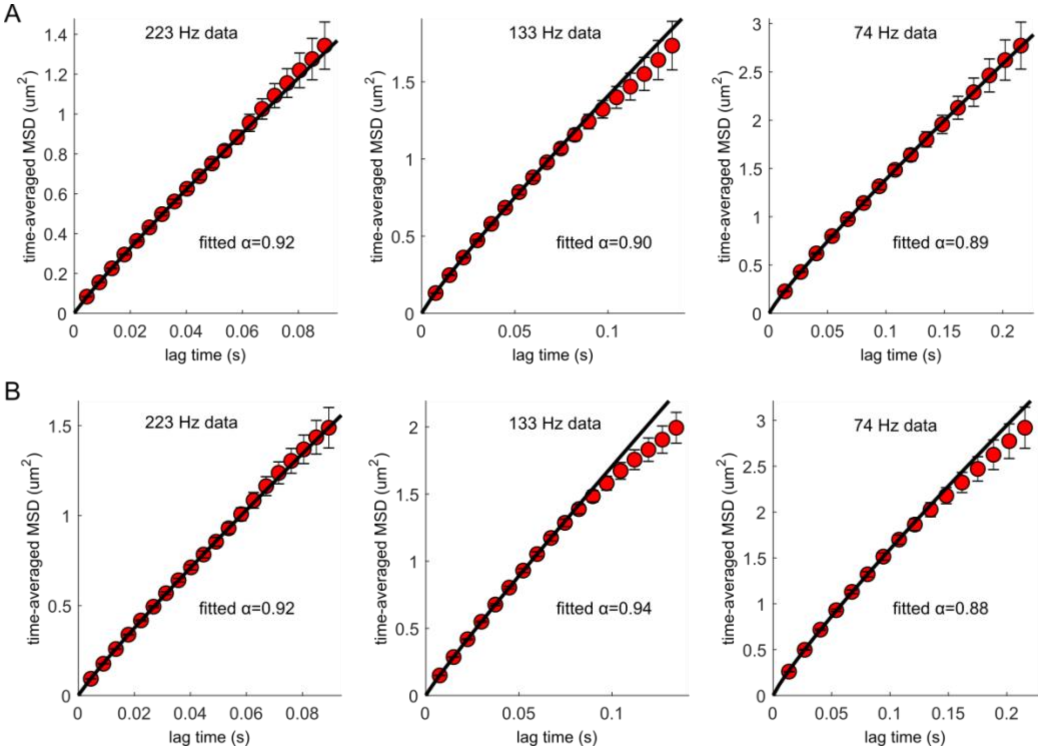


**Figure 2-Figure Supplement 2. MSD from model simulations**. The MSD was estimated form the Brownian simulation, using the same algorithm described at the Material and Methods and describe at Supplementary Figure 1. The MSD and anomalous exponent correspond to the data shown for the data plotted in Figure 2D-H. (**A**) High binding probability $P_{trap} = 0.99$ and $k_{on} = 1000$. (**B**) Smaller binding probability $P_{trap} = 0.1$ and $k_{on} = 500$.

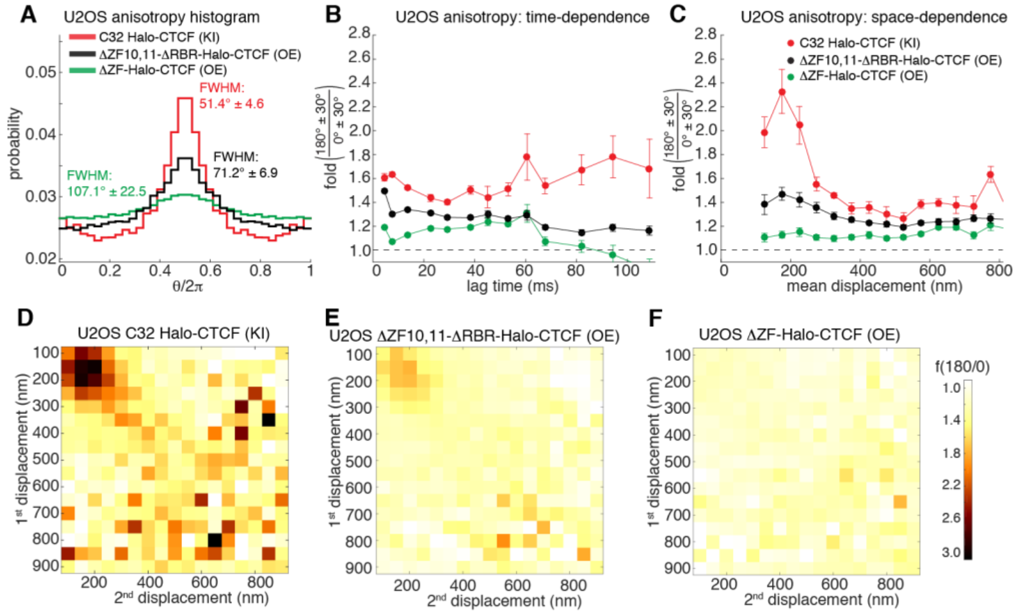


**Figure 3-Figure Supplement 1. Anisotropy wild-type and mutant CTCF in human U2OS cells.**
Anisotropy plots for C32 Halo-CTCF (knock-in), Halo-CTCF without Zinc Finger 10 and 11 and the RBR (transiently expressed; ΔZF10,11-ΔRBR-Halo-CTCF) and Halo-CTCF without all 11 Zinc Fingers (transiently expressed; ΔZF-Halo-CTCF) in human U2OS cells showing fold-anisotropy, $f_{180/0}$, at the bulk level (**A**), as a function of the lag time (**B**) and as a function of the mean displacement length (**C**). (**D-F**) Anisotropy heatmaps. Plot of $f_{180/0}$ as a function of the length of the first and second displacement for the indicated cell lines.

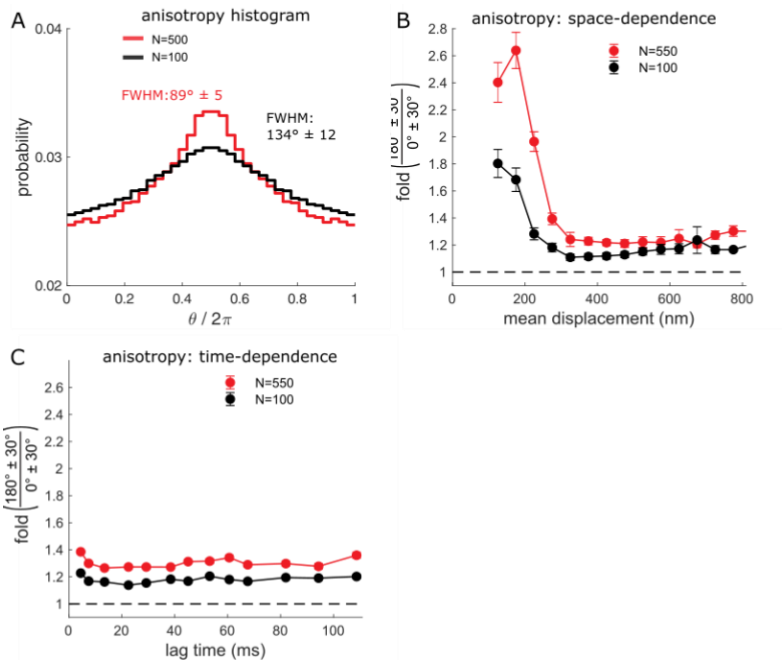


**Figure 3-Figure Supplement 2. Effect of TTZ abundance on anisotropy.** (**A**) Angle distribution of the trajectory computed from the simulation when the nucleus contains only zones of type 1 and 2 (CBSs and TTZs). The nucleus contains a high number of trapping zones (N=500) (red curve). The nucleus contains a small number of trapping zones (N=100) (black curve). (**B**) Plot of f180/0 vs. mean displacement length averaging over all lag times. (**C**) Plot of f180/0 vs. lag time averaging over all displacement lengths.

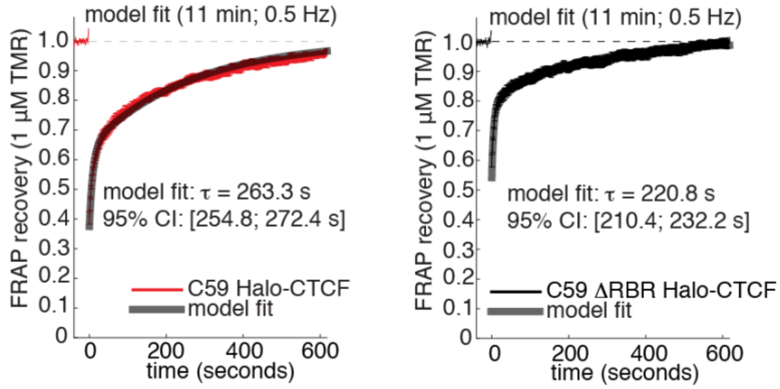


**Figure 4-Figure Supplement 1. Model-fit to FRAP data.**
Raw FRAP data (330 frames at 1 frame every 2 seconds and bleach applied at frame 20) for mESC C59 Halo-CTCF (left) and mESC C59D2 ΔRBR-Halo-CTCF (right). A 2-state reaction dominant model was fit and the slow-component interpreted as the residence time for binding to cognate DNA binding sites (shown as well as 95% confidence interval).

#### Video Legends

**Video 1**. Single Halo-CTCF protein exhibiting anomalous diffusion inside mESC nucleus.
Frame-rate: 134 Hz; dye: PA-$JF_{549}$; laser excitation pulse: 1 ms of 561 nm; Cell line: C59. Raw microscopy data with tracking overlaid in red.

**Video 2**. Single Halo-CTCF protein exhibiting anomalous diffusion inside mESC nucleus.
Frame-rate: 134 Hz; dye: PA-$JF_{549}$; laser excitation pulse: 1 ms of 561 nm; Cell line: C59. Raw microscopy data with tracking overlaid in red.

**Video 3**. Single Halo-CTCF protein exhibiting anomalous diffusion inside mESC nucleus.
Frame-rate: 134 Hz; dye: PA-$JF_{549}$; laser excitation pulse: 1 ms of 561 nm; Cell line: C59. Raw microscopy data with tracking overlaid in red.

**Video 4**. Single Halo-CTCF protein exhibiting anomalous diffusion inside mESC nucleus.

Frame-rate: 134 Hz; dye: PA-$JF_{549}$; laser excitation pulse: 1 ms of 561 nm; Cell line: C59. Raw microscopy data with tracking overlaid in red.

**Table S1. Parameters for the simulation**

| Description | Parameters | Value |
|---|---|---|
| Polymer persistence length | $l_0$ | 600nm |
| Diffusion constant of the free protein | $D_F$ | $5\mu m^2/\text{sec}$ |
| Diffusion constant of the protein inside a zone (TTZ or PTZ) | $D_Z$ | $2.5\mu m^2/\text{sec}$ |
| Diffusion constant of the protein inside a CBS | $D_{CBS}$ | $0.5\mu m^2/\text{sec}$ |
| Trapping radius of CBS/TTZ | $\varepsilon_{CBS}\ /\varepsilon_{TTZ}$ | 30nm/200nm |
| Release radius of the protein from CBS/TTZ | $a_{CBS}\ /a_{TTZ}$ | 40nm/210nm |
| Characteristic time bound at the cognate binding site. Randomized from a Poisson distribution with mean $\tau_{\text{CBS}}$ | $\tau_{\text{CBS}}$ | 1 minutes |
| Spring constant | $k$ | $\frac{3k_BT}{(0.2l_0)^2}Nm^{-1}$ |
| Monomer-monomer LJ distance | $\sigma$ | $l_0$ |
| Radius of the nucleus | $A$ | $5\mu\text{m}$ |
| Fraction of cognate binding sites | $f_{CBS}$ | 0.02 |
| Fraction of the Power-law-distributed-Trapping Zones out total zone number | $f_{PT}$ | (Figure4H-I): 0.2 (red,black) |
| Monomer number/ total number of zones | $N$ | (Figure2D-H): 500<br>(Figure4H-I): 500 (red), 500 (black), 550 (green), 500 (purple) |
| Size distribution of the Power-law Trapping Zones | $P_{PTZ}(\varepsilon)\sim A_2\varepsilon^{-\gamma}$ | (Figure4H-I): $\gamma = 0.5$ |

| | | |
|---|---|---|
| (PTZ) | | |
| Release radius of the protein from the PTZ. Different for each PTZ, depending on its size CBS/TTZ | $a_{PTZ}$ | $a_{PTZ} = \varepsilon_{PTZ} + 10\text{nm}$ |
| Trapping probability to the Transiently Trapping Zone (TTZ) | $P_{trap,TTZ}$ | (Figure2D-H): 0.99 (red), 0.1 (black)<br>(Figure4H-I): 0.99 (red), 0.2 (black), 0.1 (green), 0.99 (purple) |
| Trapping probability to the cognate binding site (CBS) | $P_{trap,CBS}$ | Equal to $P_{trap,TTZ}$ |
| Trapping probability to the Power-law Trapping Zone (PTZ) | $P_{trap,PTZ}$ | The trapping probability in a PDZ is given by<br>$P_{trap,PTZ} = \frac{A_1}{\varepsilon^{\delta}}$<br>$A_1$ is a constant equal to 100 |
| Trapping exponent of the PTZs $P_{trap,PTZ}(\varepsilon) \sim \varepsilon^{-\delta}$ | $\delta$ | (Figure4H-I): 1.2 (red, black) |
| Exit probability when hitting the boundary of a zone | $P_{exit}$ | (Figure2D-H): 0.3<br>(Figure4H-I): 0.3 (red), (0.3) |
| On-Rate to bind inside a TTZ or a PTZ | $k_{on}$ | (Figure2D-H): 1msec$^{-1}$ (red), 0.5msec$^{-1}$ (black) |
| Off-Rate to be released from a binding site inside a TTZ | $k_{off}$ | 10000 msec$^{-1}$ |
